# Type I PRMTs and PRMT5 Inversely Regulate Post-Transcriptional Intron Detention

**DOI:** 10.1101/2021.08.20.457069

**Authors:** Maxim I. Maron, Alyssa D. Casill, Varun Gupta, Simone Sidoli, Charles C. Query, Matthew J. Gamble, David Shechter

## Abstract

Protein arginine methyltransferases (PRMTs) are required for the regulation of RNA processing factors. Type I enzymes catalyze mono- and asymmetric dimethylation; Type II enzymes catalyze mono- and symmetric dimethylation. To understand the specific mechanisms of PRMT activity in splicing regulation, we inhibited Type I and II PRMTs and probed their transcriptomic consequences. Using the newly developed SKaTER-seq method, analysis of co-transcriptional splicing revealed that PRMT inhibition resulted in slower splicing rates. Surprisingly, altered co-transcriptional splicing kinetics correlated poorly with ultimate changes in alternative splicing of polyadenylated RNA—particularly intron retention (RI). Investigation of RI following inhibition of nascent transcription demonstrated that PRMTs inversely regulate RI post-transcriptionally. Subsequent proteomic analysis of chromatin-associated polyadenylated RNA identified aberrant binding of the Type I substrate, CHTOP, and the Type II substrate, SmB. Targeted mutagenesis of all methylarginine sites in SmD3, SmB, and SmD1 recapitulated splicing changes seen with Type II PRMT inhibition. Conversely, mutagenesis of all methylarginine sites in CHTOP recapitulated the splicing changes seen with Type I PRMT inhibition. Closer examination of subcellular fractions indicated that RI were isolated to the nucleoplasm and chromatin. Together, these data demonstrate that PRMTs regulate the post-transcriptional processing of nuclear, detained introns through Sm and CHTOP arginine methylation.

## INTRODUCTION

The mammalian genome encodes nine protein arginine methyltransferases (PRMTs 1-9; PRMT4 is also known as CARM1). Arginine methylation is critical in regulating signal transduction, gene expression, and splicing (reviewed by Guccione and Richard 2019; Lorton and Shechter 2019). For instance, an important function of PRMT5 along with its cofactors pICln and MEP50 (also known as WDR77), is assembly of small nuclear ribonucleoproteins (snRNPs)— core components of the spliceosomal machinery (Meister et al. 2001; Neuenkirchen et al. 2015). This includes both non-enzymatic chaperoning of Sm proteins via PRMT5/pICln following their translation and post-translational methylation of SmD1 (SNRPD1), SmD3 (SNRPD3), and SmB/B’ (SNRPB), by PRMT5-MEP50 (Friesen et al. 2001; Meister et al. 2001; Paknia et al. 2016). Following their methylation, these Sm proteins are delivered to SMN where they are bound to small nuclear RNAs (snRNAs) in preparation for further processing and eventual nuclear import (Boisvert et al. 2002; Meister and Fischer 2002; Pellizzoni et al. 2002). Disruption of PRMT5 leads to numerous splicing defects, primarily exon skipping (SE) and intron retention (RI) (Boisvert et al. 2002; Bezzi et al. 2013; Koh et al. 2015; Braun et al. 2017; Fedoriw et al. 2019; Fong et al. 2019; Radzisheuskaya et al. 2019; Tan et al. 2019; Li et al. 2021; Sachamitr et al. 2021). Intron retention is a highly prevalent alternative splicing event in tumors and is ubiquitous across cancer types (Dvinge and Bradley 2015). Therefore, understanding the mechanisms that govern RI and the connection to PRMTs is of great interest to further advance our understanding of this disease.

RNA splicing can occur during transcriptional elongation or after the transcript has been cleaved and released from polymerase (reviewed by Neugebauer 2019). Although the importance of PRMT5 in preserving splicing integrity is clear, whether PRMT5 exerts its influence over co- or post-transcriptional splicing is still unknown. Previous work has indirectly implicated PRMT5 in the regulation of post-transcriptional splicing in *Arabidopsis* and as a regulator of detained introns (DI)—unspliced introns in poly(A) RNA that remain nuclear and are removed prior to cytoplasmic export (Braun et al. 2017; Jia et al. 2020). DI are highly conserved, enriched in RNA processing factors, and their removal has been proposed as part of a feedback mechanism to control protein expression during differentiation and cell stress (Yap et al. 2012; Wong et al. 2013; Braunschweig et al. 2014; Boutz et al. 2015; Pimentel et al. 2016). Sm proteins have been implicated in regulating DI as PRMT5 inhibition leads to loss of Sm methylation and a simultaneous increase in DI (Braun et al. 2017). Nonetheless, a direct connection of DI or RI levels to Sm protein function and arginine methylation has yet to be demonstrated.

The mechanistic role of Type I PRMTs (PRMT1-4, 6, 8) in splicing is still poorly understood. Recent reports demonstrate that there are consequences on SE following Type I PRMT inhibition, with many of the proteins altered in methylation having a potential role in RNA post-transcriptional regulation (Fedoriw et al. 2019; Fong et al. 2019). Consistent with a role in post-transcriptional processing, the primary Type I methyltransferase—PRMT1—has been implicated in the regulation of RNA export through methylation of the Transcription and Export (TREX) components—Aly and REF export factor (ALYREF) and chromatin target of PRMT1 (CHTOP) (Hung et al. 2010; Van Dijk et al. 2010; Chang et al. 2013). Loss of CHTOP leads to an accumulation of nuclear polyadenylated transcripts due to a block in RNA export (Chang et al. 2013). Furthermore, ALYREF and CHTOP are both deposited co-transcriptionally along RNA and are bound to RI (Viphakone et al. 2019). As the nuclear export of RNA is intimately linked to splicing—dissociation of splicing factors is a prerequisite for successful export—regulation of this process is an attractive means to control RI (Luo and Reed 1999). Despite this, the role of nuclear export factors and the impact of Type I PRMTs on regulating RI remains unexplored.

Here, we test both the co- and post-transcriptional consequences of PRMT5 or Type I PRMT inhibition (PRMTi). We also examine which factors are responsible for PRMTi-mediated splicing consequences. As both Type I PRMTs and PRMT5 have been extensively reported to be required for the pathogenicity of lung cancer (Hwang et al. 2021)—yet their role in the transcription and splicing of this disease remains largely unstudied—we used A549 human alveolar adenocarcinoma cells as our model. Using GSK591 (also known as EPZ015866 or GSK3203591), a potent and selective inhibitor of PRMT5 (Duncan et al. 2016), and MS023, a potent pan-Type I inhibitor (Eram et al. 2016), we demonstrate that PRMTs inversely regulate RI post-transcriptionally through Sm and CHTOP arginine methylation.

## RESULTS

### Type I PRMT or PRMT5 inhibition promotes changes in alternative splicing

PRMTs consume S-adenosyl methionine (SAM) and produce S-adenosyl homocysteine (SAH) to catalyze the post-translational methylation of either one or both terminal nitrogen atoms of the guanidino group of arginine (**Figure 1a**) (Gary and Clarke 1998). All PRMTs can generate monomethyl arginine (Rme1). Type I PRMTs further catalyze the formation of asymmetric N^G^,N^G^-dimethylarginine (Rme2a); Type II PRMTs (PRMT5 and 9) form symmetric N^G^,N^’G^-dimethylarginine (Rme2s). PRMT5 is the primary Type II methyltransferase (Yang et al. 2015). As previous reports indicated that lengthy treatment with PRMTi promotes aberrant RNA splicing, we wanted to determine whether alternative splicing differences with PRMTi occurred as early as day two and, if so, how they evolved over time (Bezzi et al. 2013; Fong et al. 2019; Radzisheuskaya et al. 2019; Tan et al. 2019; Li et al. 2021; Sachamitr et al. 2021). Therefore, we performed poly(A)-RNA sequencing on A549 cells treated with DMSO, GSK591, MS023, or both inhibitors in combination for two-, four-, and seven-days. Sequencing at seven days for cotreatment was not performed as this resulted in significant cell death (not shown). Using replicate Multivariate Analysis of Transcript Splicing (rMATS) (Shen et al. 2014) to identify alternative splicing events, at day two we observed significant differences in RI with PRMTi (FDR < 0.05) relative to DMSO (**Figure 1b**). Furthermore, whereas RI were increased with GSK591- and co-treatment—signified by a positive difference in percent spliced in (+ΔPSI or +ΔΨ)—RI were decreased with MS023 treatment (-ΔΨ) relative to DMSO.

**Figure 1.**
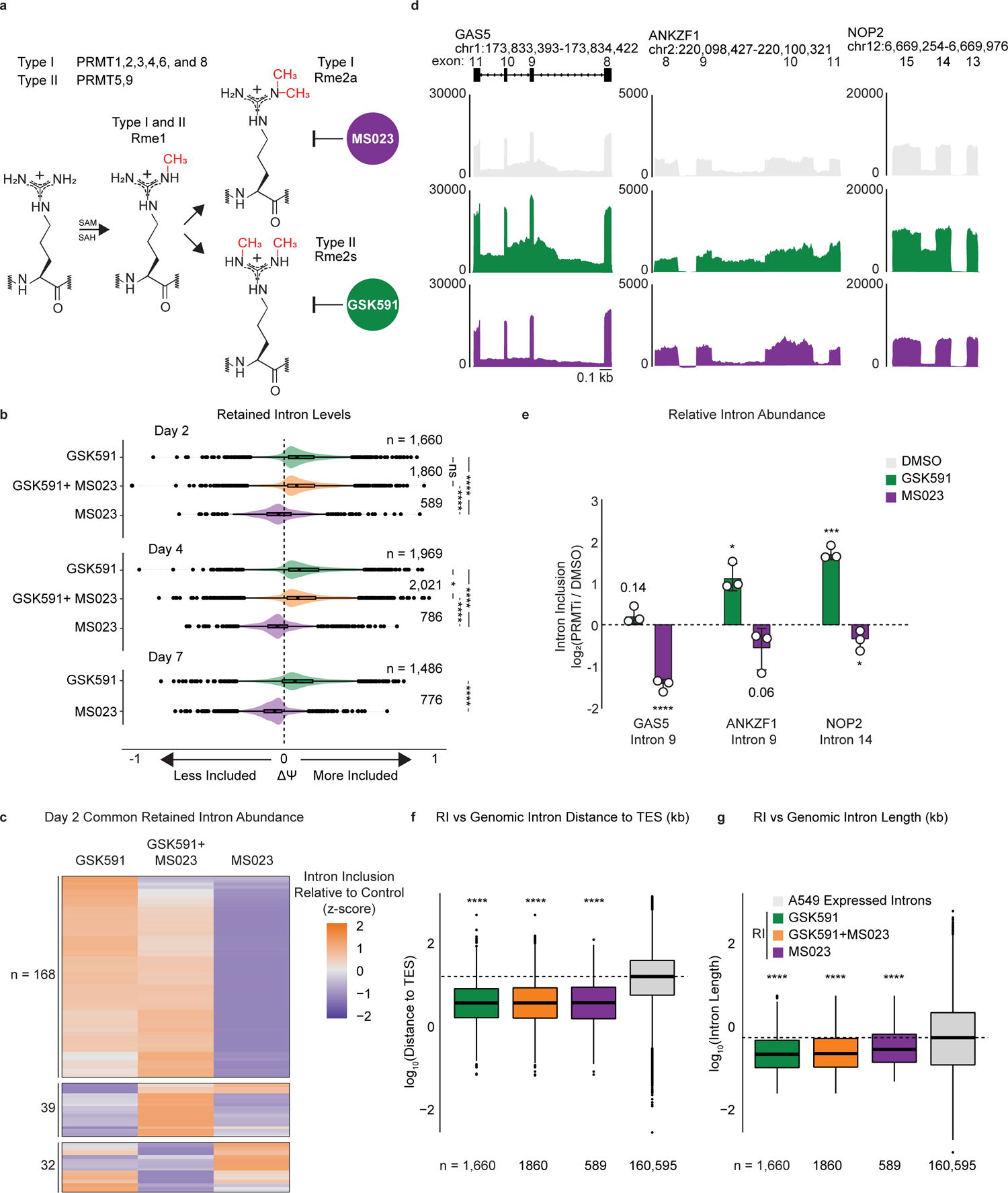
Type I PRMTs and PRMT5 inversely regulate intron retention. **a.** Overview of protein arginine methyltransferases and their catalyzed reactions. **b.** Comparison of ΔΨ for retained introns (RI) following PRMTi where ΔΨ = Ψ (PRMTi) – Ψ (DMSO). Significance determined using Kolmogorov-Smirnov test; * < 0.05, **** < 0.0001, ns = not significant. **c.** Comparison of ΔΨ z-score for common RI after two-day treatment with PRMTi. **d.** Genome browser track of poly(A)-RNA seq aligned reads for *GAS5*, *ANKZF1*, and *NOP2*. Yellow shading denotes RI. Scale (0.1 kb) indicated in lower right corner. **e.** RT-qPCR of RI highlighted in panel d. Data are represented as mean ± SD. Significance determined using Student’s t-test; * < 0.05, ** < 0.01, *** < 0.001, **** < 0.0001, ns = not significant. **f-g.** Comparison of RI and A549-expressed intron distance to the transcription-end site (TES) **(f)** or intron length **(g)** in log_10_(kb). Dashed line indicates genomic median; solid line within boxplot is condition-specific median. Significance determined using Wilcoxon rank-sum test; **** < 0.0001.

We next intersected all common RI between treatment conditions in A549 at day two (**Figure 1c**). We found 239 common introns. Strikingly, whereas GSK591 and co-treatment with GSK591 and MS023 had increased inclusion, for the same introns MS023 alone resulted in decreased inclusion relative to DMSO (**Figure 1c**). To confirm and quantify the presence of PRMT-regulated RI by RT-qPCR, we selected three genes, *GAS5*, *ANKZF1*, and *NOP2.* These were identified by rMATS as containing inversely regulated RI by either PRMT5 or Type I PRMTi (**Figure 1d**). Notably, while the expression of *GAS5* and *NOP2* was increased with GSK591, it was unchanged with MS023. Using *GAS5* intron 9, *ANKZF1* Intron 9, and *NOP2* Intron 14 primers that spanned the intron-exon boundaries and normalized by exonic primers, at day two following PRMTi we confirmed the presence of the RI (**Figure 1e**). Consistent with our rMATS data, *GAS5* intron 9, *ANKZF1* intron 9, and *NOP2* intron 14 had increased retention upon GSK591 treatment (*P* = 0.141, 0.016, 0.001, respectively). Furthermore, despite decreased dynamic range inherent to quantifying intronic signal loss, we observed less retention with MS023 treatment (*P* < 0.0001, 0.064, 0.040, respectively) (**Figure 1e**).

To confirm that these RI were specific to PRMTi and not a consequence of off-target effects of GSK591 or MS023 we generated A549 cells expressing dCas9-KRAB-MeCP2 (Yeo et al. 2018). Independent transduction of two unique guide RNAs (gRNA) for four days targeting either the major Type I methyltransferases—PRMT1 and PRMT4—or the Type II methyltransferase, PRMT5 achieved efficient knockdown (**Suppl. Figure 1a**). Furthermore, we recapitulated increased RI in *GAS5* intron 9, *ANKZF1* intron 9, and *NOP2* intron 14 by RT-qPCR following knockdown of PRMT5 with both unique gRNAs (**Suppl. Figure 1b**). We also observed reduced RI in *GAS5* intron 9, *ANKZF1* intron 9, but not *NOP2* intron 14 when analyzing RI following transduction with the stronger PRMT1 gRNA (**Suppl. Figure 1b**). Knockdown of PRMT4 only reduced RI in *GAS5* intron 9 with one gRNA, but not in *ANKZF1* intron 9 or *NOP2* intron 14. Together, these data support that the RI seen following Type I or PRMT5i are specific, yet the consequences of MS023 treatment on RI are likely more robust than CRISPR interference due to the inhibition of multiple Type I enzymes and the ability of PRMTs to scavenge each other’s substrates (Dhar et al. 2013; Eram et al. 2016).

### PRMTi-mediated RI are conserved across cell types and share common characteristics

We next asked whether the RI in our data were common to other datasets in which PRMT activity was perturbed. To accomplish this, we used rMATS on publicly available data (Braun et al. 2017; Fedoriw et al. 2019; Fong et al. 2019; Radzisheuskaya et al. 2019). In these experiments–conducted in a variety of cell lines from diseases including acute myeloid leukemia (THP-1), chronic myeloid leukemia (K562), pancreatic adenocarcinoma (PANC03.27), and glioblastoma (U87)–arginine methylation was inhibited via PRMT knockdown using CRISPRi or with PRMTi. As demonstrated by the high odds ratio (log_2_(OR) > 6) between all the datasets, we showed that there was a highly significant (Fisher’s exact adjusted *P* ≤ 1e-05) overlap in RI (**Suppl. Figure 2a**). This further supports that the consequences of PRMTi on RI are not off-target effects of the inhibitors and are also common to multiple cell types.

RI have previously been reported to have common characteristics such as being shorter, closer to transcription end site (TES), and having reduced splice site strength (Braunschweig et al. 2014). To determine the common characteristics of the PRMT-regulated RI, we analyzed their length, distances to the TES, and sequences. We found that the RI were significantly closer to the TES and shorter when compared to the genomic distribution of A549 expressed introns (*P* < 2.2e-16) (**Figure 1f and 1g**). Moreover, in analyzing the probability of nucleotide distribution at the 5’ and 3’ splice sites we noted both a preference for guanine three nucleotides downstream of the 5’ splice site and increased frequency of cytosine in the polypyrimidine tract (**Suppl. Figure 2b**). This is consistent with previous literature demonstrating that RI have common features contributing to their persistence in poly(A) RNA (Bezzi et al. 2013; Braunschweig et al. 2014; Braun et al. 2017; Tan et al. 2019).

### PRMT-dependent changes in co-transcriptional splicing do not reflect changes in mRNA

The inverse relationship on RI by Type I or PRMT5i in addition to their TES-proximal location, shorter length, and weaker splice sites led us to investigate the kinetics of co-transcriptional splicing. To accomplish this, we used Splicing Kinetics and Transcript Elongation Rates by Sequencing (SKaTER seq) (Casill et al. 2021). As splicing changes are present as early as two days following GSK591 or MS023 treatment, we used this time point of PRMTi for our analysis. Briefly, SKaTER seq uses a three-hour 5,6-dichloro-1-β-D-ribofuranosylbenzimidazole (DRB) treatment to synchronize transcription, followed by a rapid wash-out to allow productive elongation to commence. Once RNA pol II begins elongating, starting at 10 minutes nascent RNA is collected every five minutes until 35 minutes post-DRB washout (**Figure 2a**). Nascent RNA isolated via a 1 M urea wash of chromatin and an additional poly(A)-RNA depletion is then sequenced. The rate of nascent RNA formation—including: (1) RNA pol II initiation and pause-release (spawn) rate, (2) elongation rate, (3) splicing rate, and (4) transcript cleavage rate—is then calculated by using a comprehensive model that determines the rates that best fit the sequencing coverage (Casill et al. 2021).

**Figure 2.**
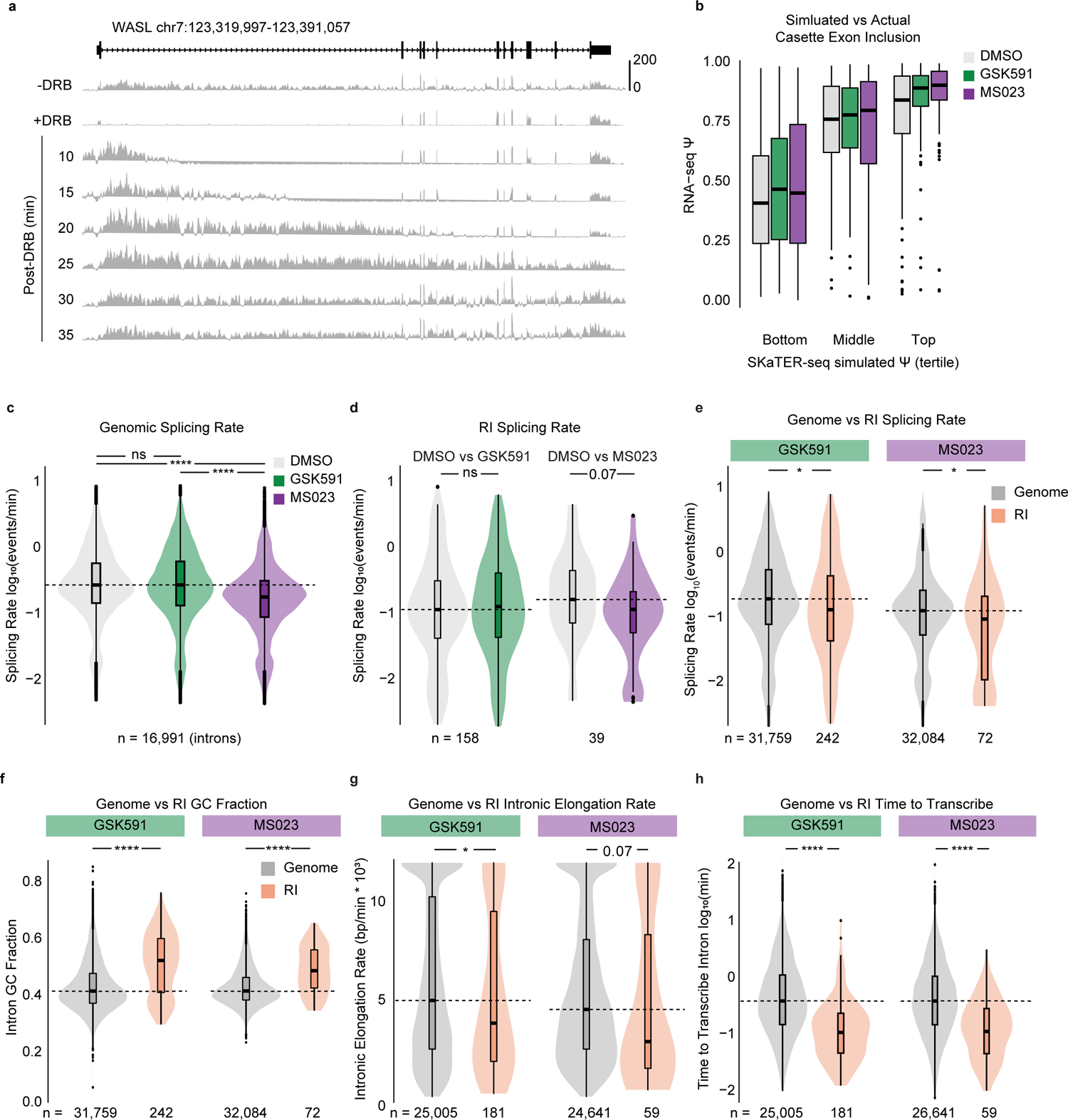
RI share unique characteristics and are independent of PRMT-regulated co-transcriptional splicing. **a.** Histogram of SKaTER-seq aligned reads across *WASL*. **b.** Correlation of SKaTER-seq model simulated cassette exon Ψ versus poly(A)-RNA seq Ψ. **c.** Distribution of splicing rates for common genomic introns. Dashed line indicates DMSO median; solid line within boxplot is condition-specific median. Significance determined using Wilcoxon rank-sum test; **** < 0.0001, ns = not significant. **d.** Distribution of splicing rates for common retained introns as in panel c. **e-h.** Distribution of splicing rates **(e)**, GC fraction **(f)**, intronic elongation rate **(g)**, and time to transcribe **(h)** for RI (orange) and A549 expressed introns (dark gray) within the same condition. Dashed line indicates genomic median; solid line within boxplot is condition-specific median. Significance determined using Wilcoxon rank-sum test; * < 0.05, **** < 0.0001.

To assess the accuracy of the rates determined by the SKaTER model, we used the spawn, elongation, splicing, and cleavage rates to simulate a predicted poly(A)-RNA cassette exon Ψ and compared these results with our poly(A)-RNA sequencing. We successfully predicted cassette exon Ψ detected in poly(A) RNA in DMSO (Spearman’s correlation coefficient (ρ) = 0.54), GSK591 (ρ = 0.61), and MS023 (ρ = 0.55) (*P* < 2.2e-16) treatment conditions (**Figure 2b**). Next, we compared spawn rate to poly(A)-RNA sequencing transcripts per million (TPM). As predicted, we observed a strongly significant (*P* < 2.2e-16) correlation between RNA pol II spawn rate and TPM in DMSO-, GSK591-, and MS023-treated cells (ρ = 0.50, 0.51, 0.51, respectively) (**Suppl. Figure 3a**) (Casill et al. 2021). Consistent with previously published reports, a comparison of splicing rates within each condition confirmed that constitutive introns splice faster than cassette exons as well as alternative 5’ and 3’ splice sites (**Suppl. Figure 3b**) (Pandya-Jones et al. 2013).

As poly(A)-RNA sequencing revealed opposing alternative splicing changes—specifically increased RI with GSK591 and decreased RI with MS023—we next asked how PRMTi affected the global distribution of splicing rates relative to DMSO. Surprisingly, GSK591 treatment did not significantly change the median of this distribution (Wilcoxon rank-sum test, *P* > 0.05), while MS023 treatment resulted in significantly slower global splicing rates relative to DMSO (*P* < 2.2e-16) (**Figure 2c**). As our primary interest was in RI, we compared the splicing rates of introns that were retained with PRMTi to the same introns in DMSO-treated cells. We found no significant difference in the median of GSK591-treated cells compared to DMSO (*P* = 0.53), while MS023 treatment similarly led to slower splicing rates (*P =* 0.07) (**Figure 2d**). This result was inconsistent with the changes seen in poly(A) RNA as a slower splicing rate should increase RI. Therefore, the co-transcriptional splicing rate suggested that changes in RI following PRMTi were not due to co-transcriptional splicing.

### RI share unique characteristics that limit their co-transcriptional removal

We next asked how RI splicing rates compared to non-RI. We observed that introns that tended to be retained in GSK591- or MS023-treated cells were slower to splice (*P* = 0.01 and *P* = 0.04, respectively) when compared to the genomic distribution (**Figure 2e**). As we previously found that RI were more likely to be closer to the TES and had slower splicing rates (**Figure 1f and 1g**), we asked whether intron position correlated with splicing rate. Indeed, we determined that introns located closer to the TES had slower splicing rates (**Suppl. Figure 4a**). When examining additional characteristics of RI, we noted that they had a significantly higher GC fraction in both GSK591-(0.52, *P* = 1.63e-15) and MS023-treated cells (0.50, *P* = 3.89e-6) relative to the global median (0.41) (**Figure 2f**). As GC content influences polymerase elongation, we analyzed the intronic elongation rate of RI in comparison to non-RI. Consistent with an increased GC content, we found that RI had slower elongation rates relative to non-RI (*P* = 0.07 and 0.02 for GSK591 and MS023, respectively) (**Figure 2g**). Next, as previous reports implicated polymerase elongation rate with co-transcriptional splicing outcomes—splice site availability dictates splicing decisions—we wanted to understand if the slower elongation rate seen with RI compensated for their shorter size (Fong et al. 2014). To accomplish this, we assessed how long RNA pol II took to transcribe RI compared to non-RI. We found that despite their slower elongation rate, RI transcription was completed significantly faster compared to non-RI in both GSK591-and MS023-treated cells (*P* = 6.22e-17 and *P* = 8.12e-6, respectively) (**Figure 2h**). Given the proximity of RI to the TES, this likely further decreased their ability to be removed co-transcriptionally. Taken together, these results support the conclusion that RI in GSK591- and MS023treated cells share unique characteristics that increase their probability of post-transcriptional processing.

### PRMTs regulate RI post-transcriptionally

The paradox that RI are decreased with Type I PRMTi—despite their slower co-transcriptional splicing—led us to hypothesize that PRMTs exert their control over splicing post-transcriptionally. To test this hypothesis, we used the elongation, splicing, and cleavage rates determined by SKaTER seq to calculate the probability that a transcript will be cleaved from RNA pol II prior to a given intron being spliced. Consistent with most splicing being co-transcriptional, we found that the median probability of cleavage prior to splicing was 9.7% in DMSO-treated cells and 8.7% in GSK591-treated cells, yet the probability with MS023 treatment was significantly higher at 18.1% (**Figure 3a**). The global distribution was significantly reduced with GSK591 treatment (*P* = 0.002) and increased with MS023 treatment (*P* < 2.2e-16) when compared to DMSO (**Figure 3a**). We next analyzed the probability of cleavage prior to splicing for RI. Consistent with the hypothesis that PRMTs regulate splicing of RI post-transcriptionally, the median probability of transcript cleavage prior to splicing was significantly higher for RI (50.6% and 63.2%, respectively) when compared to non-RI (11.7% and 23.1%, respectively; *P* = 5.74e-10 and 0.0002) (**Figure 3b**). Furthermore, intron position was strongly predictive of whether transcript cleavage was likely to occur prior to splicing: TES proximal introns had a higher probability of cleavage prior to their being spliced (**Suppl. Figure 4b**). All told, the proximity of RI to the TES, their shorter length, and their slower splicing rate likely drives their decreased probability of being spliced co-transcriptionally.

**Figure 3.**
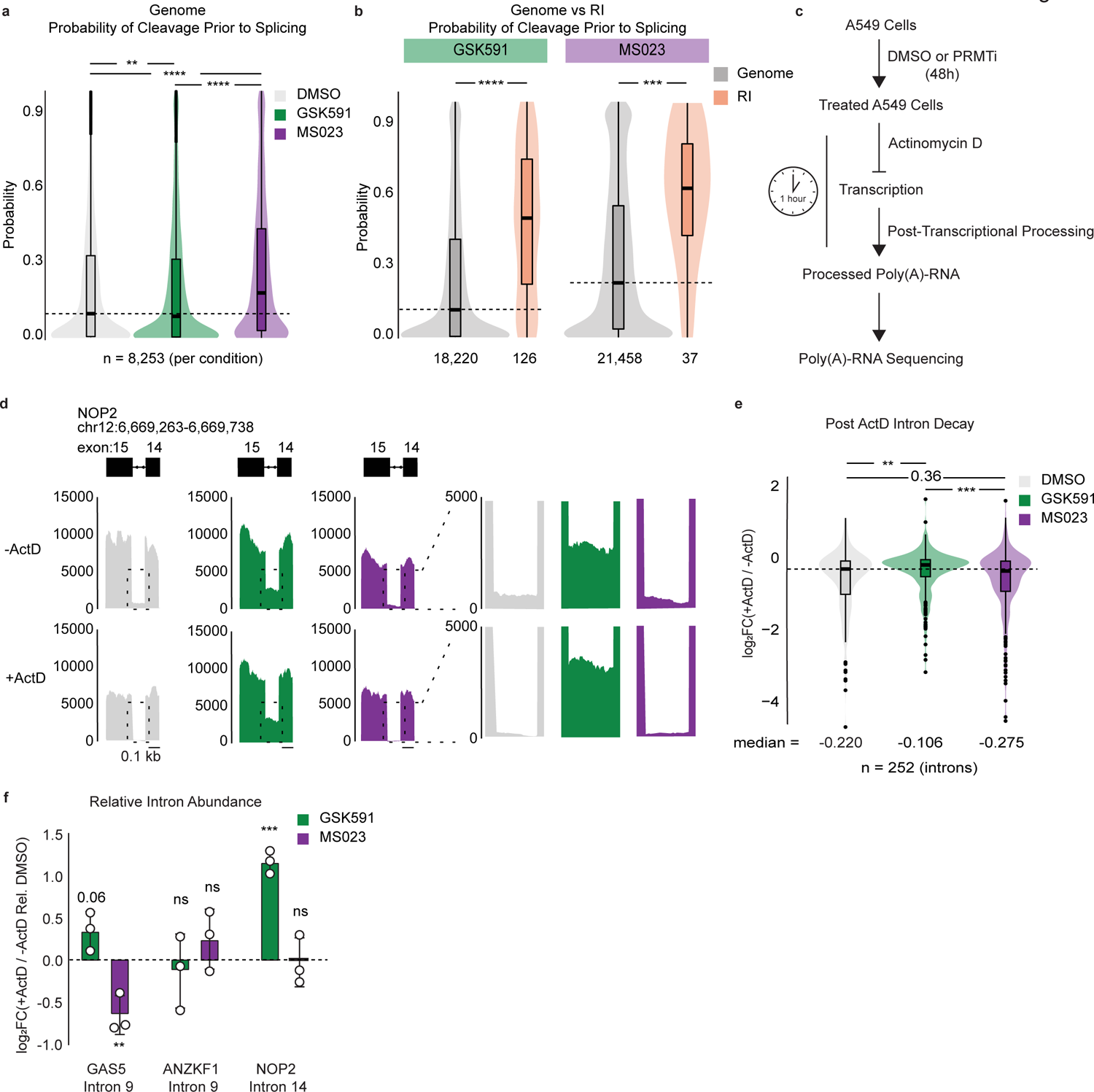
Type I PRMTs and PRMT5 regulate RI post-transcriptionally. **a.** Probability of cleavage prior to splicing for common introns relative to DMSO. Dashed line indicates DMSO median; solid line within boxplot is condition-specific median. Significance determined using Wilcoxon rank-sum test; ** < 0.01, **** < 0.0001. **b.** Probability of cleavage prior to splicing for RI (orange) and A549 expressed introns (dark gray) within the same condition. Dashed line indicates genomic median; solid line within boxplot is condition-specific median. Significance determined using Wilcoxon rank-sum test; *** < 0.001, **** < 0.0001. **c.** Overview of Actinomycin D (ActD) poly(A)-RNA sequencing protocol. **d.** Genome browser track of poly(A)-RNA seq aligned reads (-) and (+) ActD for *NOP2* **e.** Distribution of log_2_(+ActD/-ActD) abundance for common RI between GSK591 and MS023 relative to DMSO. Dashed line indicates DMSO median; solid line within boxplot is condition-specific median. Significance determined using Wilcoxon rank-sum test; ** < 0.01, *** < 0.001. **f.** RT-qPCR of RI (-) and (+) ActD relative to DMSO. Data are represented as mean ± SD. Significance determined using Student’s t-test; ** < 0.01, *** < 0.001, ns = not significant.

The slowing of co-transcriptional splicing following PRMTi suggests that there would be an increase in observed RI. Paradoxically, poly(A)-RNA sequencing revealed that although RI are increased in GSK591-treated cells, they are reduced in MS023-treated cells. This led us to hypothesize that MS023 promotes more efficient post-transcriptional intron loss, whereas the opposite is true for GSK591. To test this model, we pre-treated A549 cells for two days with GSK591 or MS023 and then blocked transcription using actinomycin D (ActD) for 60 minutes. We then performed poly(A)-RNA sequencing to determine the post-transcriptional consequences of PRMTi on RI (**Figure 3c**). Genome browser tracks demonstrated reduced RI and flanking exon signal intensity following 60 minutes of ActD treatment (**Figure 3d**). Further quantification of RI common to both GSK591- and MS023-treated cells relative to their -ActD controls—normalized to the abundance of their flanking exons—demonstrated a negative median log_2_ fold change in all conditions, indicating less RI abundance in the ActD-treated samples. Moreover, consistent with a post-transcriptional mechanism, we observed significantly less intron loss in GSK591-treated cells (median = −0.106) compared to DMSO-(−0.220) or MS023-treated cells (−0.275) (*P =* 0.002 relative to DMSO) (**Figure 3e**). The RI loss following ActD in MS023-treated cells was not significantly different than DMSO, although the median was more negative suggesting a trend toward increased decay (*P* = 0.36) (**Figure 3e**).

We next validated these changes in RI following treatment with ActD using RT-qPCR for our three candidate introns described above (**Figure 1e**). GSK591 treatment resulted in significantly slower intron decay relative to DMSO for *GAS5* intron 9 and *NOP2* intron 14 but not *ANKZF1* intron 9 after 60 minutes of ActD treatment (*P* = 0.06, 0.0005, 0.74, respectively) (**Figure 3f**).

Conversely, intron decay was faster with MS023 treatment relative to DMSO for *GAS5* intron 9 but not *ANKZF1* intron 9 or *NOP2* intron 14 (*P* = 0.0043, 0.27, 0.98, respectively) (**Figure 3f**). As the changes in RI following transcriptional inhibition with ActD reflect those of PRMTi alone— increased RI with GSK591 and decreased RI with MS023—these results support that PRMTs regulate RI post-transcriptionally.

### PRMTi alters binding of RNA processing factors to chromatin-associated poly(A) RNA

We next sought to identify the factors responsible for mediating the post-transcriptional consequences of PRMTi. Previous reports have highlighted that delayed co-transcriptional splicing leads to chromatin retention of poly(A) transcripts (Brody et al. 2011; Pandya-Jones et al. 2013; Yeom et al. 2021). Therefore, we analyzed differences in proteins bound to nuclear, and more specifically, chromatin-associated poly(A) RNA. To accomplish this, we performed UV-crosslinking of A549 cells two days after treatment with PRMTi. We then isolated the chromatin fraction, and after a high-salt wash, sheared the material with a light, brief sonication. This was followed by poly(A) enrichment using oligo(dT) beads. To control for non-specific interactions, we performed high stringency washes and added an excess of free poly(A) to our negative control. After elution of the bound poly(A) RNA, we digested the RNA and analyzed the remaining material with liquid chromatography coupled online with tandem mass spectrometry (LC-MS/MS) (**Figure 4a**).

**Figure 4.**
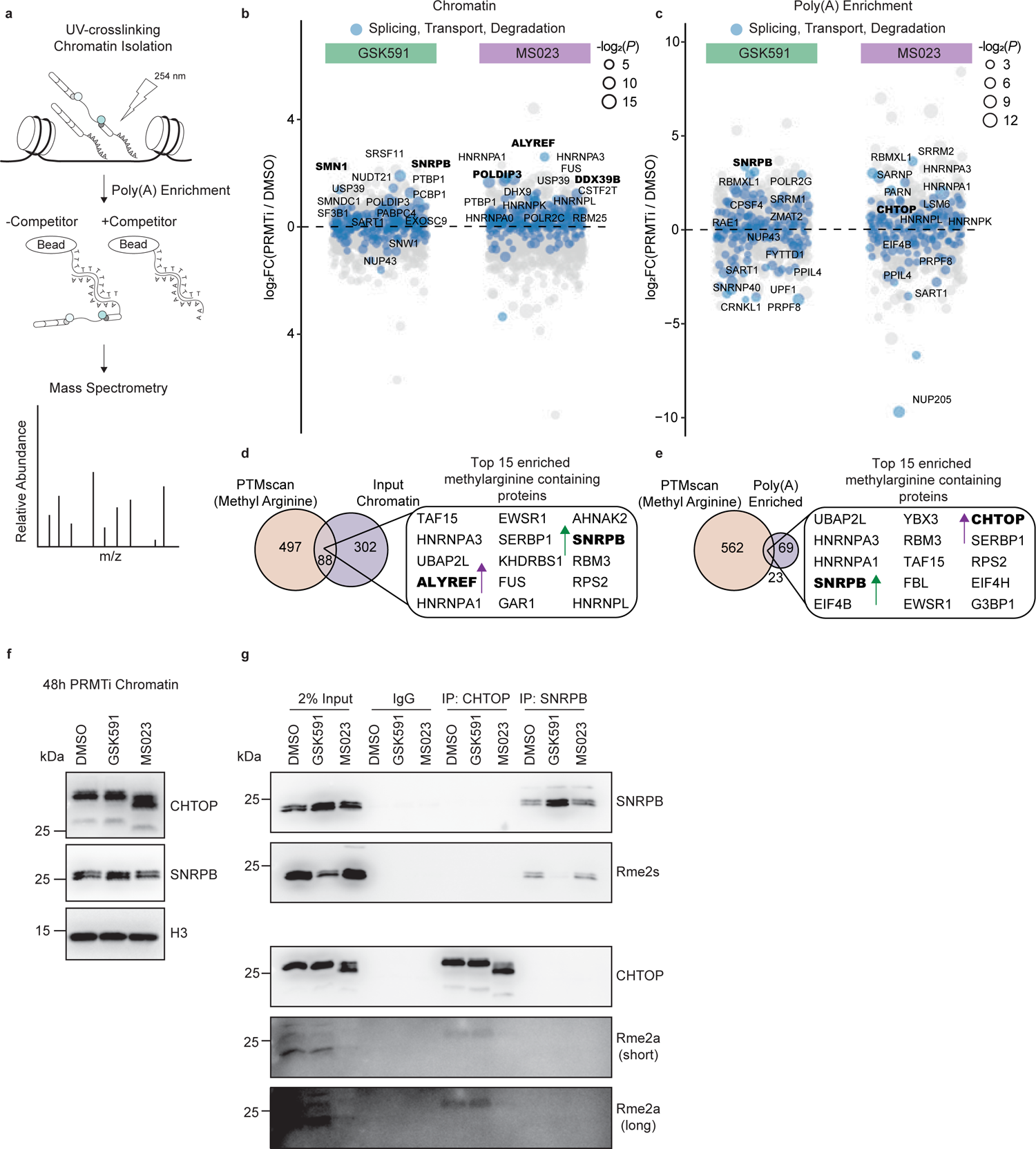
PRMTi promotes aberrant binding of RNA processing factors to chromatin-associated poly(A) RNA. **a.** Overview of chromatin-associated poly(A)-RNA LC-MS/MS experiment. **b-c.** Dot plot of proteins bound to chromatin **(b)** or chromatin-associated poly(A) RNA **(c)** relative to DMSO. Circle size is proportional to -log_2_(*P*). Colored values denote factors with ontology pertaining to RNA splicing, transport, or degradation. The names of the top 15 significant proteins are labeled. Significance determined using a heteroscedastic t-test. **d-e.** Venn diagram comparing proteins containing methylarginine (PTMscan) and those that were differentially enriched in the chromatin **(d)** or chromatin-associated poly(A) **(e)** fractions following PRMTi. **f.** Western blot of chromatin following two-day treatment with DMSO, GSK591, or MS023. **g.** Immunoprecipitation and analysis of CHTOP and SNRPB methylarginine following treatment with DMSO, GSK591, or MS023 for two days.

We identified 1,832 unique proteins in the input chromatin across the three conditions (**Suppl. Table 1**). Following GSK591 treatment, 118 had significantly altered abundance with 70 increased and 48 decreased (*P* < 0.05). In the MS023-treated input chromatin, we observed 255 differentially enriched proteins with 150 increasing in abundance and 105 decreasing (*P* < 0.05). The poly(A) enriched fraction—after background subtraction—contained 1,251 unique proteins. Of these, 32 were differentially bound in the GSK591-treated samples with 24 increased and 8 decreased (*P* < 0.05). In the MS023-treated cells, there were 55 proteins with altered binding— 37 increased and 18 decreased (*P* < 0.05).

To identify the biological processes most affected by these inhibitors, we performed over representation analysis of the top 200 enriched proteins **(Suppl. Figures 5a and 5b**). In the chromatin fraction of both GSK591- and MS023-treated cells—consistent with the gross aberrations in splicing following treatment with these inhibitors—we observed a significant enrichment of gene ontology terms for RNA splicing and ribonucleoprotein complex biogenesis (*Padj* < 0.05) (**Suppl. Figure 5a**). Interestingly, in GSK591-treated cells, there was also a unique enrichment for the regulation of protein localization to Cajal bodies (*Padj* < 0.05). Conversely, in MS023-treated cells we observed a highly significant enrichment for nuclear RNA export and the regulation of RNA catabolism (*Padj* < 0.05). When looking specifically at the poly(A)-enriched fraction, we noted similar themes as those in the input chromatin (**Suppl. Figure 5b**).

**Figure 5.**
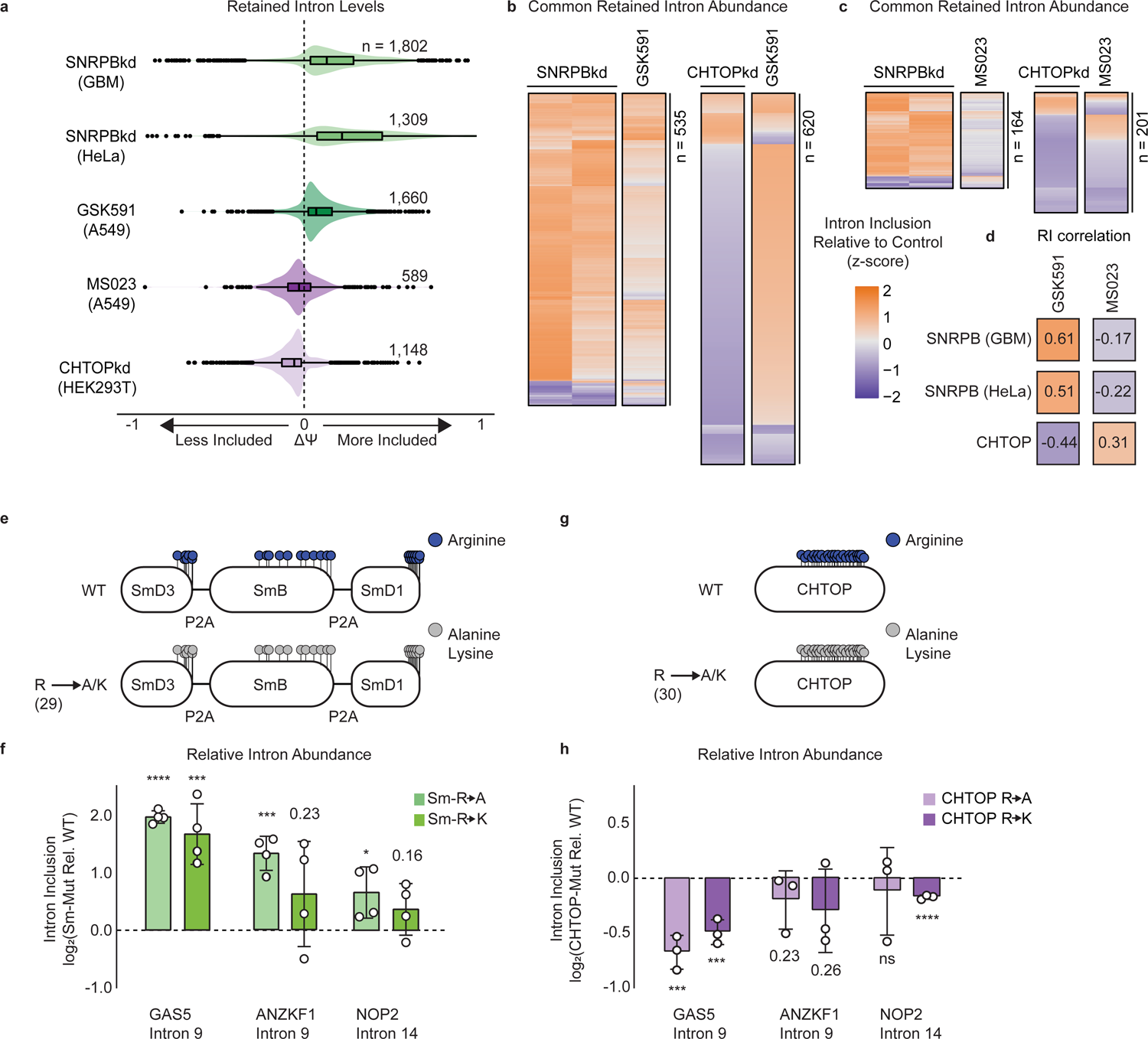
Sm and CHTOP methylarginine mediates PRMT-dependent RI. **a.** Comparison of ΔΨ for RI following SNRPB knockdown (kd), GSK591, MS023, and CHTOPkd where ΔΨ = Ψ (Treatment) – Ψ (Control). **b-c.** Comparison of ΔΨ z-score for common RI between SNRPBkd and CHTOPkd with either GSK591 **(b)** or MS023 **(c)**. **d.** Spearman rank correlation of ΔΨ for common RI between SNRPBkd and CHTOPkd with either GSK591 or MS023. **e.** Schematic of Sm expression constructs where lollipops represent individual methylarginines. **f.** RT-qPCR of RI following transduction with Sm R-to-A or R-to-K mutants relative to wildtype (WT). Data are represented as mean ± SD. Significance determined using Student’s t-test; * < 0.05, *** < 0.001, **** < 0.0001. **g.** Schematic of CHTOP expression construct as in panel e. **h.** RT-qPCR of RI following transduction with CHTOP R-to-A or R-to-K mutants relative to WT. Data are represented as mean ± SD. Significance determined using Student’s t-test; *** < 0.001, **** < 0.0001, ns = not significant.

### PRMTi perturbs binding of nuclear export factors and snRNPs to poly(A) RNA

Of the proteins that were differentially bound to the input chromatin and poly(A) RNA following treatment with PRMTi, we were interested in candidates that were most likely to mediate post-transcriptional RI. To accomplish this, we took into consideration the enriched gene ontology categories described above and highlighted the most significant factors involved in RNA splicing, transport, and degradation relative to their log_2_ fold change (**Figure 4b and 4c**). In the input chromatin of GSK591-treated cells, we observed increased signal in the snRNP chaperone, SMN, as well as the snRNP component, SNRPB (SmB/B’) (**Figure 4b**). We also observed enrichment of SNRPB in the poly(A)-enriched fraction (**Figure 4c**). Previous reports have documented changes in exon skipping following SNRPB knockdown—including within its own transcript—that resemble changes seen with PRMT5 knockdown (Saltzman et al. 2011; Bezzi et al. 2013). Together with the fundamental role of snRNPs in splicing, this suggested that SNRPB may also be involved in the regulation of GSK591-mediated RI.

In the input chromatin of MS023-treated cells we observed an increase of the TREX components ALYREF, DDX39B, and POLDIP3 relative to DMSO (**Figure 4b**). ALYREF and DDX39B are recruited to pre-mRNA co-transcriptionally and are crucial for the eventual hand-over of the mRNA to the nuclear export receptor, nuclear RNA export factor 1 (NXF1) (Heath et al. 2016). When looking specifically at the poly(A) RNA fraction of MS023-treated cells, we noted an increase in CHTOP, also a component of the TREX complex (**Figure 4c**) (Heath et al. 2016). Both CHTOP and ALYREF have been shown to be direct targets of PRMT1—the major Type I methyltransferase and target of MS023 (Hung et al. 2010; Van Dijk et al. 2010). In both cases, arginine methylation of ALYREF and CHTOP was demonstrated to weaken their binding to RNA and promote interaction with NXF1—consistent with their increase in both the input chromatin and poly(A) RNA fractions following MS023-mediated inhibition of Type I PRMT activity (Hung et al. 2010; Chang et al. 2013).

To further refine the factors likely to be responsible for PRMTi-dependent RI, we compared these data with proteins that contained methylarginine in our PTMScan methylome data (Maron et al. 2021). When we intersected the arginine methylome with the differentially enriched proteins in our input and poly(A) enriched fractions (*P* < 0.1), there were 88 and 23 unique methylarginine containing proteins, respectively. We then analyzed the top 15 differentially enriched proteins in each fraction. In the input, we noted that ALYREF and SNRPB were both modified with methylarginine (**Figure 4d**). Furthermore, when looking at the poly(A) fraction, we again observed SNRPB as well as CHTOP (**Figure 4e**). The prior identification of ALYREF, CHTOP, and SNRPB as methylarginine containing proteins suggested that these *bona fide* PRMT substrates may be potential mediators of PRMT-dependent RI (Brahms et al. 2001; Hung et al. 2010; Van Dijk et al. 2010; Maron et al. 2021).

To validate changes in CHTOP and SNRPB chromatin-association following PRMTi, we isolated the chromatin fraction and probed for CHTOP and SNRPB. Consistent with abundant GR repeats—the preferred substrate motif for PRMT1 and PRMT5—present in CHTOP, we observed faster CHTOP gel migration consistent with hypomethylation in MS023-treated cells (**Figure 4f**). Furthermore, in line with the proteomic analysis of input chromatin following GSK591 treatment, there was increased signal intensity of SNRPB (**Figure 4f**). To further confirm the changes in methylarginine on CHTOP and SNRPB, we performed an immunoprecipitation for these proteins following PRMTi (**Figure 4g**). Subsequent analysis of methylarginine confirmed loss of CHTOP Rme2a with MS023 treatment and SNRPB Rme2s with GSK591 treatment. Taken together, the differential enrichment of ALYREF, DDX39B, and POLDIP3 on chromatin, along with CHTOP on poly(A) RNA following treatment with MS023—in addition to their dependence on Rme2a—presented the possibility that the TREX complex may have an important role in the post-transcriptional consequences of Type I PRMTs on RI. Moreover, as snRNPs are fundamental in splicing, increased SNRPB in both the input and poly(A) fractions following treatment with GSK591—as well as the loss of Rme2s—prompted us to further evaluate the role of SNRPB in the regulation of RI.

### Knockdown of CHTOP and SNRPB recapitulates PRMTi-mediated RI

To address the question of whether CHTOP, ALYREF, or SNRPB were involved in regulation of RI, we first checked whether there was any publicly available RNA-seq data in which these proteins were perturbed. We found two independent datasets where SNRPB was knocked down in U251 glioblastoma cells or HeLa cells (Saltzman et al. 2011; Correa et al. 2016). We also identified two independent datasets where CHTOP or ALYREF were knocked down in HEK293T cells or HeLa cells, respectively (Fan et al. 2019; Viphakone et al. 2019). Strikingly, after performing rMATS on these datasets, we observed that both SNRPB knockdown (SNRPBkd) experiments strongly recapitulated the increase in RI seen with GSK591 treatment (**Figure 5a**). Conversely, although ALYREF knockdown did not significantly affect RI levels (**Suppl. Figure 6a**), CHTOP knockdown (CHTOPkd) resulted in a global decrease in RI inclusion paralleling that seen with MS023 treatment (**Figure 5a**). To understand if the RI were common across datasets, we intersected either the SNRPBkd or CHTOPkd RI with PRMTi-dependent RI (**Figures 5b and 5c**). There were 535 mutual RI between GSK591 and both the SNRPBkd datasets, most of which had the same effect on ΔΨ (log_2_ Odds Ratio 7.84 for GBM and 7.56 for HeLa, *P <* 1e-300) (**Figure 5b**). The opposite was true when comparing GSK591 to CHTOPkd—we observed 620 shared RI (log_2_ Odds Ratio 7.49, *P* < 1e-300) that largely contrasted in their ΔΨ (**Figure 5b**). MS023 treatment had the inverse effect of GSK591 when compared to SNRPBkd and CHTOPkd. There were 164 overlapping RI between both SNRPBkd datasets (log_2_ Odds Ratio 6.68 for GBM and 6.38 for HeLa, *P <* 7e-290) with the majority having opposite ΔΨ values; CHTOPkd and MS023 treatment shared 201 RI (log_2_ Odds Ratio 646, *P <* 2e-279), with the majority having coincident ΔΨ values (**Figure 5c**). Consistent with PRMTs regulating a common group of RI, SNRPBkd and CHTOPkd were significantly correlated with either GSK591 or MS023 treatment (**Figure 5d**). Remarkably, the correlations that were positive for GSK591 treatment were negative for MS023 treatment, reflecting the changes seen in poly(A)-RNA seq and consistent with SNRPB and CHTOP mediating the effects of PRMTi on RI.

### Sm arginine mutants increase RI

To address the question of whether Sm arginine methylation was directly involved in the regulation of RI, we designed a single vector containing SNRPD3, SNRPB, and SNRPD1—the Sm targets of PRMT5—as either wildtype, R-to-A, or R-to-K mutants separated by 2a-self cleaving peptides (P2A) and modified with FLAG-(SNRPD3 and SNRPB) or V5-affinity (SNRPD1) tags at their N-termini (**Figure 5e**). To identify the appropriate arginines to mutate, we utilized our PTMScan data as well as a previously published high-resolution crystal structure of the U4/U6.U5 tri-snRNP (Charenton et al. 2019; Maron et al. 2021). As we recently showed that PRMTs prefer to methylate intrinsically disordered regions (IDRs), we mutated all 29 arginines within Sm IDRs and within GR repeats (**Figure 5e**) (Maron et al. 2021). We then transduced A549 cells with the constructs and performed RT-qPCR for *GAS5* intron 9, *ANKZF1* intron 9, or *NOP2* intron 14. We achieved similar expression of the exogenous Sm proteins relative to the endogenous between wildtype and mutant constructs at the level of RNA (**Suppl. Figure 6b**). However, likely owing to the meticulous control of total Sm levels by the cell, we were unable to achieve ample overexpression with the wildtype vector (**Suppl. Figure 6c**) (Prusty et al. 2017). We observed a migratory shift toward a lower molecular weight in the R-to-A mutants when compared to the wildtype or R-to-K mutants (**Suppl. Figure 6c**). We also observed a strong Rme2s signal on the FLAG-SmB wildtype protein that was absent on the R-to-A and R-to-K mutants (**Suppl. Figure 6c**). Furthermore, and consistent with a role for Sm methylarginine in regulating RI levels, following Sm R-to-A mutagenesis, we observed increased inclusion in all three RI candidates (*P* ≤ 0.05) (**Figure 5f**). We also detected a significant increase in *GAS5* intron 9 (*P* < 0.001) with the Sm R-to-K mutants and a trend toward increased RI in *ANKZF1* intron 9 (*P* = 0.23) and *NOP2* intron 14 (*P* = 0.16). Thus, mutagenesis of Sm methylarginine sites increased RI in our candidate transcripts, yet the greater effect of the R-to-A mutants suggests that the charge of arginine itself may also play an important role in regulating RI levels.

### CHTOP arginine mutants decrease RI

We performed similar experiments with CHTOP whereby we mutated 30 arginines present within the centrally located “GAR” motif—the preferred PRMT1 substrate recognition motif—to either alanine or lysine (**Figure 5g**). This region of CHTOP has been previously shown to be required for PRMT1-catalyzed methylarginine (Van Dijk et al. 2010). We transduced A549 cells with the constructs and performed RT-qPCR for *GAS5* intron 9, *ANKZF1* intron 9, or *NOP2* intron 14. We achieved similar expression of the mutant CHTOP proteins when compared to the wildtype at the level of RNA and protein (**Suppl. Figure 6c and 6d**). Interestingly, when transducing cells with the wildtype CHTOP, we noted a gross downregulation of the endogenous transcript (**Suppl. Figure 6e**). CHTOP has been previously reported to control its own expression as part of an autoregulatory loop (Izumikawa et al. 2016). We did not observe this compensation with either the R-to-A or R-to-K mutants. Consistent with the gel shift in CHTOP seen following MS023 treatment, we observed a similar change in the R-to-A mutant when compared to the wildtype or R-to-K mutants. Moreover, supporting a role for CHTOP methylarginine in Type I PRMT-dependent RI inclusion, in both R-to-A and R-to-K mutants we observed decreased inclusion in *GAS5* intron 9 (*P* < 0.001) (**Figure 5h**). We also saw a trend toward decreased inclusion of *ANKZF1* intron 9 (*P* = 0.23 and 0.26) and significantly decreased inclusion of *NOP2* intron 14 (*P* < 0.0001) in the R-to-K mutant, but not the R-to-A mutant. Taken together, these experiments support that CHTOP methylarginine is involved in regulating RI levels.

**Figure 6.**
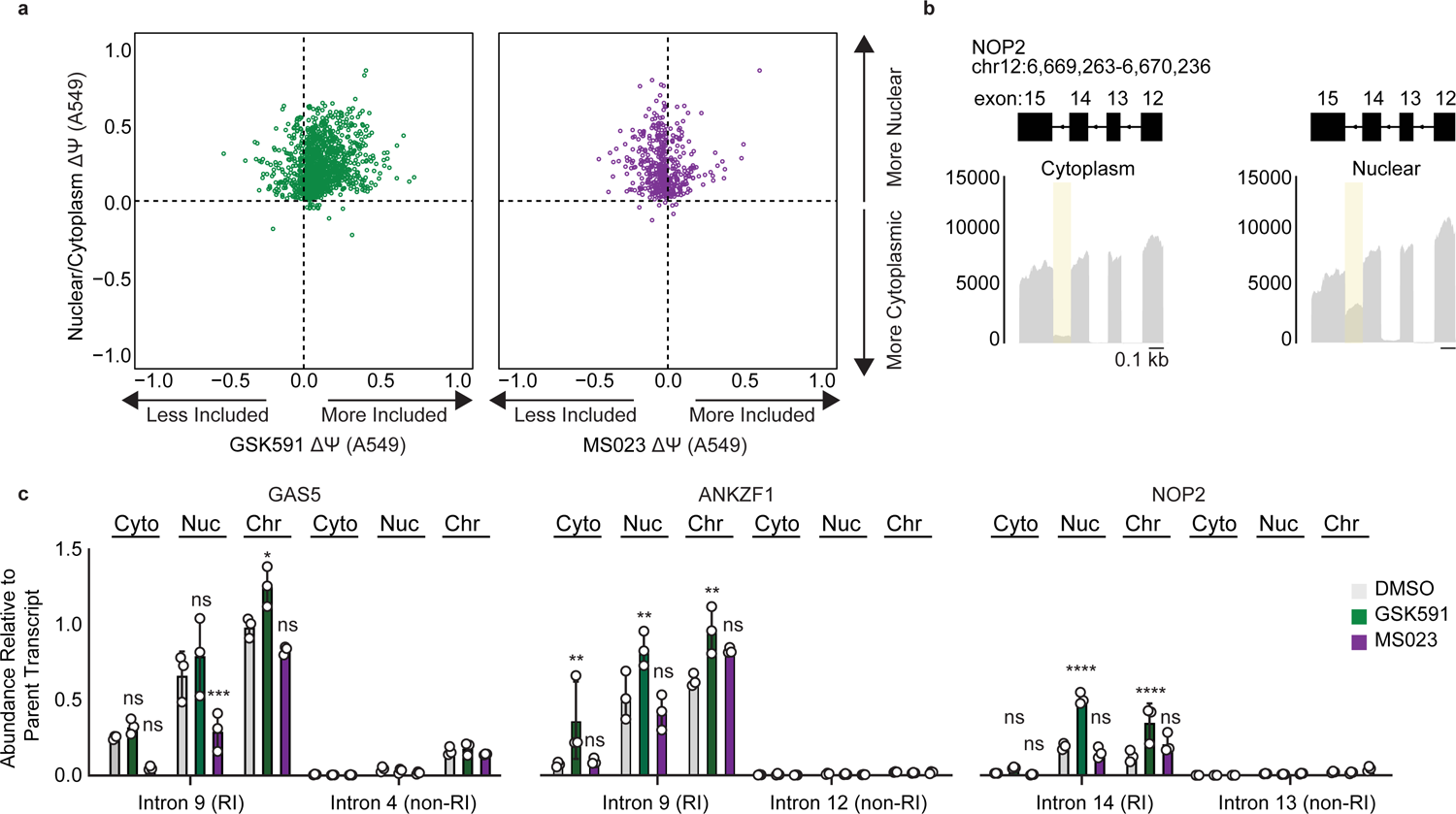
PRMT-dependent RI are localized to the nucleoplasm and chromatin. **a.** Scatter plot of ΔΨ for common RI in A549 nuclear/cytoplasmic fractions and GSK591 (left, green) or MS023 (right, purple) where ΔΨ = Ψ (Nuclear/PRMTi) – Ψ (Cytoplasm/DMSO). **b.** Genome browser track of poly(A)-RNA seq aligned reads for *NOP2* in A549 cytoplasmic or nuclear fractions. **c.** RT-qPCR of RI or non-RI from cytoplasmic, nuclear, or chromatin fractions of A549 cells treated with DMSO, GSK591, or MS023 for two days. Data are represented as mean ± SD. Significance determined using two-way analysis of variance with Tukey’s multiple comparisons test; * < 0.05, ** < 0.01, *** < 0.001, **** < 0.0001, ns = not significant.

### PRMT-regulated RI are detained within the nucleoplasm and chromatin

PRMT5 has been proposed to specifically regulate DI—introns that persist in poly(A) RNA but remain nuclear (Braun et al. 2017). However, direct experimental evidence—namely fractionation of subcellular compartments to identify the location of RI—has been lacking. To address the question of whether the RI in our data are nuclear and therefore DI, we first ran rMATS on publicly available ENCODE poly(A)-RNA seq from cytoplasmic and nuclear fractions of A549 cells (ENCSR000CTL and ENCSR000CTM, respectively). We observed an enrichment of RI within the nuclear fraction—97% of significant RI events (FDR < 0.05) had a +ΔΨ, where ΔΨ is the difference in Ψ between the nuclear and cytoplasmic compartments. We then intersected these data with the PRMTi-mediated RI and noted that there was a significant overlap with both GSK591-(log_2_ OR 8.89, *P* < 1e-300) and MS023-treatment (log_2_ OR 7.75, *P* < 1e-300) when compared to all A549 expressed introns (**Figure 6a**). We also observed that *GAS5* intron 9, *ANKZF1* intron 9, and *NOP2* Intron 14 were significantly increased in nuclear poly(A) RNA (*GAS5* and *ANKZF1* not shown) (**Figure 6b**).

To validate that these RI remain nuclear following PRMTi, and to further resolve their localization within the nucleus, we treated A549 cells with PRMTi and isolated cytoplasmic, nucleoplasmic, and chromatin fractions. We then quantified levels of *GAS5* intron 9, *ANKZF1* intron 9, and *NOP2* intron 14 or non-RI within the same poly(A) transcripts (**Figure 6c**). In accordance with the RI being localized to the nucleus, in DMSO-, GSK591-, and MS023-treated cells, we observed a significant increase in nucleoplasmic and chromatin signal compared to the cytoplasm (*P* < 0.05) (**Figure 6c**). Importantly, there was no significant difference between cytoplasmic or nuclear enrichment of the non-RI tested. Together, these data support that these RI are indeed localized to the nucleoplasm and chromatin fractions and are thus DI. Therefore, we conclude that Type I PRMTs and PRMT5 regulate post-transcriptional DI through modulation of snRNP and CHTOP arginine methylation (**Figure 7a**). We also propose the following model for PRMT-dependent DI regulation: (1) snRNP Rme2s is required for post-transcriptional maturation of DI-containing transcripts—in the absence of Rme2s these snRNPs are bound to transcript but the DI are sequestered from splicing or degradation (**Figure 7b**), (2) CHTOP Rme2a is necessary for DI-containing transcripts to progress through the TREX-mediated nuclear export pathway; CHTOP lacking Rme2a inhibits this progression but does not protect the mRNA, which together with the consequent increase in nuclear residence time promotes splicing or degradation (**Figure 7c**).

**Figure 7.**
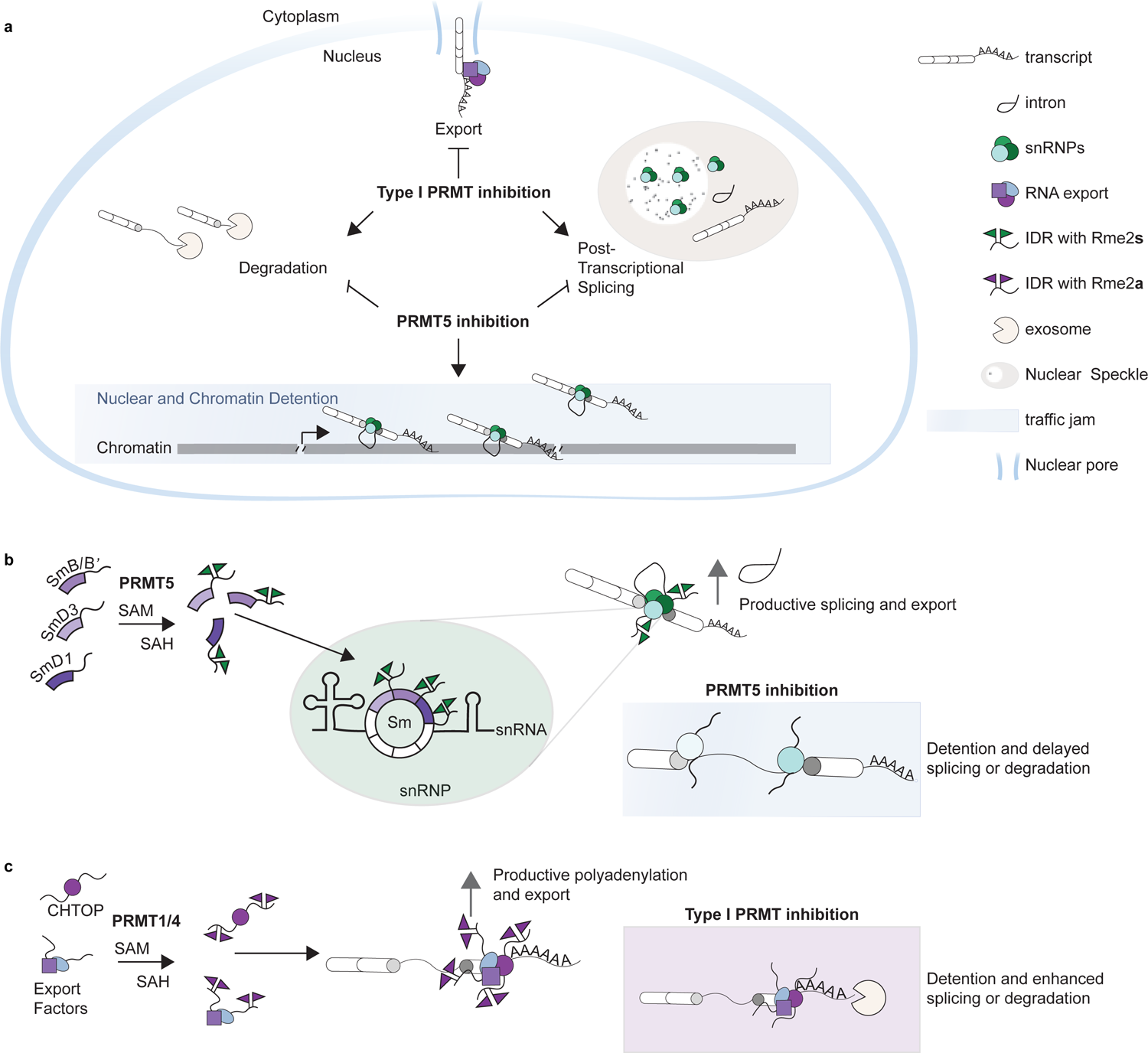
Type I PRMTs and PRMT5 regulate post-transcriptional intron detention through Sm and CHTOP arginine methylation. **a-c.** Model figures representing an overview of Type I- and PRMT5-dependent regulation of detained introns (DI) **(a)**. PRMT5-catalyzed Sm Rme2s is necessary for productive splicing and maturation of DI; absence of Rme2s results in detention of the parent transcript **(b)**. Type I PRMT-catalyzed CHTOP Rme2a is required for appropriate polyadenylation and nuclear export of DI; absence of Rme2a increases nuclear residence time resulting in increased degradation or splicing **(c)**.

## DISCUSSION

In this study, our goal was to characterize the consequences of PRMTi on RI. Many reports have documented that PRMT5 has a crucial role in splicing (Boisvert et al. 2002; Bezzi et al. 2013; Koh et al. 2015; Braun et al. 2017; Fedoriw et al. 2019; Fong et al. 2019; Radzisheuskaya et al. 2019; Tan et al. 2019; Li et al. 2021; Sachamitr et al. 2021). However, the role of Type I PRMTs in splicing has only recently been appreciated (Fedoriw et al. 2019; Fong et al. 2019). This is the first study to report that Type I PRMTi decreases DI. Interestingly, in the presence of both Type I and PRMT5i, DI were increased. This suggests that PRMT5 catalyzed Rme2s is dominant in DI regulation. This is consistent with our model, whereby snRNP binding occurs upstream of CHTOP-mediated intron detention and protects RNA from post-transcriptional splicing or degradation (**Figure 7a**).

Furthermore, whether PRMTs regulate splicing co- or post-transcriptionally has not previously been explored. We found that Type I PRMTi promoted slower co-transcriptional splicing, while paradoxically resulting in less DI. While understanding the effects of Type I PRMTi on global splicing is beyond this current work, one possibility may be that given the largely negative effect arginine methylation has on protein-RNA binding (Blackwell and Ceman 2012), the absence of Rme2a may inhibit or delay dissociation of RNA processing factors from nascent RNA thereby leading to slower splicing. We and others have found that PRMT-dependent DI share characteristics such as higher GC content, non-canonical splice sites, and location closer to the TES that makes their co-transcriptional removal less likely (Wong et al. 2013; Braunschweig et al. 2014; Boutz et al. 2015; Braun et al. 2017). Consistently, when modeling co-transcriptional kinetics of DI compared to the global distribution, DI had slower splicing and elongation rates and took less time to transcribe. Together with their TES-proximal location, this combination is likely to decrease the probability that splicing will be completed co-transcriptionally. These characteristics were true for Type I PRMT inhibited cells as well—despite having less DI overall—indicating their reliance on a post-transcriptional mechanism. As has been suggested by others, DI are likely an evolutionarily conserved class of introns used for fine tuning the transcriptome (Braunschweig et al. 2014; Boutz et al. 2015; Pimentel et al. 2016). This is further supported by our comparison of DI in publicly available datasets, which—despite using diverse cell models and methods of inhibiting PRMTs—had a strong overlap with the DI present in our data. The same characteristics that lead these introns to become detained are likely what make them more sensitive to perturbation of splicing components. This could lead them to be co-opted in disease such as cancer where they have been shown to inactivate tumor suppressors and RNA export proteins (Dvinge and Bradley 2015; Jung et al. 2015).

Indeed, when using actinomycin D to study post-transcriptional DI processing, PRMT5i resulted in slower DI loss compared to DMSO or Type I PRMTi, whereas Type I PRMTi had faster DI loss relative to DMSO and PRMT5i. Thus, the post-transcriptional—rather than co-transcriptional—consequences reflect those seen in poly(A) RNA. Although little is known about how post-transcriptional splicing is accomplished, previous work has demonstrated that post-transcriptional spliceosomes are retained in nuclear speckles (Girard et al. 2012). Furthermore, CHTOP and the TREX complex have also been found to localize within nuclear speckles (Dias et al. 2010; Chang et al. 2013). Perturbation of either splicing factors or TREX components resulted in accumulation of poly(A) RNA in nuclear speckles (Dias et al. 2010; Chang et al. 2013). It is therefore possible that the two types of PRMTi coalesce at the level of this membraneless organelle. Consistent with this is the observation that SRRM1 was enriched in the poly(A) fraction in PRMT5i, whereas SRRM2 was enriched in the poly(A) fraction in Type I PRMTi (**Figure 5b**). Many SR-repeat proteins—including SRRM1 and SRRM2—are localized to nuclear speckles (Ilik et al. 2020). Given the importance of arginine methylation in regulating liquid-liquid phase separation (LLPS) and the role of LLPS in maintaining nuclear speckle integrity—as well as other membraneless organelles such as Cajal bodies—PRMTi may result in failed coordination and spatial localization of RNA processing factors leading to aberrant post-transcriptional processing (Courchaine et al. 2021; Liao and Regev 2021).

As delayed splicing has been shown to result in chromatin retention of transcripts (Brody et al. 2011; Pandya-Jones et al. 2013; Yeom et al. 2021), we assayed chromatin-associated poly(A) RNA to identify factors that regulate PRMTi-dependent DI. By interrogating changes in proteins bound to chromatin-associated poly(A) RNA—and their methylarginine levels—we identified snRNPs and the TREX complex component, CHTOP, as key-mediators of DI regulation. Interestingly, CHTOP contains a retained intron within its own transcript that is used to autoregulate its expression by nonsense-mediated decay (Izumikawa et al. 2016). This is accomplished by antagonizing intron excision by hnRNPH1 when CHTOP—using its “GAR”-motif—binds to its own transcript (Izumikawa et al. 2016). Therefore, it is possible that a similar mechanism may be occurring globally whereby introns that are slow to splice are bound by CHTOP—which has been shown to bind nascent RNA co-transcriptionally—and this prevents further binding by hnRNP proteins and intron excision (Izumikawa et al. 2016; Viphakone et al. 2019).

Another mechanism may be explained by the role of CHTOP in alternative polyadenylation (APA) (Viphakone et al. 2019). Previous work demonstrated that CHTOP specifically binds to the 3’ end of transcripts and CHTOPkd promotes changes in APA (Viphakone et al. 2019). The length of poly(A) is a driver of exosome-mediated decay, whereby abnormally long poly(A) tails—seen in detained intron containing transcripts—drive exosome-mediated decay (Bresson and Conrad 2013; Bresson et al. 2015; Muniz et al. 2015). Hyperadenylation occurs through poly(A) binding protein nuclear 1 (PABPN1) dependent stimulation of poly(A) polymerases (Bresson and Conrad 2013). In MS023-treated cells, PABPN1 was specifically enriched in the chromatin and poly(A) fraction (*P* < 0.1) (**Supplemental Table 1**). Thus, loss of CHTOP Rme2a may promote hyperadenylation and consequent exosomal degradation of DI-containing transcripts.

As PRMTs have thousands of diverse substrates—many of which are involved in RNA homeostasis—the effect of PRMTs on splicing integrity is likely to be multifactorial (Guccione and Richard 2019; Lorton and Shechter 2019). Here, we highlighted both snRNPs and CHTOP as key mediators of PRMTi-regulated DI. However, given the importance of both CHTOP and snRNPs in global RNA processing, why only a subset of alternative splicing events and transcripts are affected by their perturbation remains to be known. Importantly, there are likely many other factors involved in regulating PRMT-dependent alternative splicing. Future work will be needed to characterize the methylarginine dependent mechanisms of post-transcriptional RNA processing.

## MATERIALS AND METHODS

### Cell Culture

A549 and HEK293T cells were cultured in DMEM (Corning) supplemented with 10% FBS (Hyclone), 100 µg/mL streptomycin and 100 I.U./mL penicillin (Corning) and maintained at 37 °C with humidity and 5% CO_2_. For this study, fresh cells were purchased from ATCC and tested routinely for *Mycoplasma* (PCR, FWD primer: ACTCCTACGGGAGGCAGCAGT, REV primer: TGCACCATCTGTCACTCTGTTAACCTC).

### RT-qPCR

RNA purification was performed using RNeasy Plus (Qiagen). Isolated RNA was reverse transcribed with Moloney murine leukemia virus (MMLV) reverse transcriptase (Invitrogen) and oligo(dT) primers. LightCycler 480 Sybr Green I (Roche) master mix was used to quantitate cDNA with a LightCycler 480 (Roche). An initial 95°C for 5 minutes was followed by 45 cycles of amplification using the following settings: 95°C for 15 s, 60°C for 1 minute. Primer sequences can be found in **Supplemental Table 2**.

### Poly(A)-RNA sequencing

RNA was extracted using RNeasy^®^ Mini Kit (Qiagen) following the manufacturer’s protocol. RNA quantitation and quality control were accomplished using the Bioanalyzer 2100 (Agilent Technologies). Stranded RNA seq libraries were constructed by Novogene Genetics US. The barcoded libraries were sequenced by Novogene on an Illumina platform using 150nt paired end libraries generating ∼30-40 million reads per replicate. Reads were trimmed and aligned to the human genome (hg19) with Spliced Transcripts Alignment to a Reference (STAR) (Dobin et al. 2013). Alternative splicing events were determined using Replicate Multivariate Analysis of Transcript Splicing (rMATS, version 4.0.2) (Shen et al. 2014). Expression was determined using Kallisto (Bray et al. 2016). IGV (Broad Institute) was used as the genome browser. Graphs pertaining to RNA seq were created using Gviz (Hahne and Ivanek 2016) in R (4.0.2) and assembled in Adobe Illustrator 2020.

### Splicing Kinetics and Transcript Elongation Rates by Sequencing

SKaTER seq was performed as described (Casill et al. 2021). Briefly, A549 cells were grown with 0.01% DMSO, 1 µM GSK591 (Cayman), or 1 µM MS023 (Cayman) for two days followed by addition of 100 µM DRB (Cayman). DRB-containing media was removed, and the cells were incubated at 37 °C until the indicated time point. The cells were washed once with 4°C PBS, and lysed by addition of 1 mL CL buffer (25 mM Tris pH 7.9 at 4°C, 150 mM NaCl, 0.1 mM EDTA, 0.1% Triton X-100, 1 mM DTT) supplemented with protease inhibitor (Thermo) and *Drosophila melanogaster* S2 cell spike-in. Next, lysate was centrifuged at 845 x g for 5 minutes at 4 °C. The pellet resuspended in 1 mL CL buffer without S2 spike-in and incubated on ice for five minutes. Repeat centrifugation was performed. The supernatant was removed, and cells resuspended in 100 µL GR buffer (20 mM Tris pH 7.9, 75 mM NaCl, 0.5 mM EDTA, 50% glycerol, 0.85 mM DTT) followed by addition of 1.1 mL NL buffer (20 mM HEPES pH 7.6, 300 mM NaCl, 7.5 mM MgCl_2_, 1% NP-40, 1 mM DTT, 1 M Urea). Following a 15-minute incubation, the lysate was spun at 16,000 x g for 10 minutes and the resulting chromatin pellet was resuspended and stored in TRIzol (Thermo, 15596026) at −80 °C. RNA isolation was followed by poly(A) depletion using the NEBNext Poly(A) mRNA magnetic isolation module (NEB, E7490L). RNA quantitation and quality control were accomplished using the Bioanalyzer 2100 (Agilent Technologies). Stranded RNA-seq libraries were prepared using the KAPA RNA HyperPrep Kit with RiboErase (HMR) and KAPA Unique Dual-Indexed Adapters (Roche) according to instructions provided by the manufacturer. The barcoded paired-end libraries were sequenced by Novogene using a NovaSeq S4, generating ∼70 million reads per replicate.

### Actinomycin D post-transcriptional processing

A549 cells were grown in the presence of 0.01% DMSO, 1 µM GSK591 (Cayman), or 1 µM MS023 (Cayman) for two days. Following a two-day incubation, the media was removed and replaced with media containing 5 µg/mL actinomycin D (Sigma) with 0.01% DMSO, 1 µM GSK591, or 1 µM MS023 for 60 minutes. RNA was isolated using RNeasy^®^ Mini Kit (Qiagen) and poly(A)-RNA sequencing was performed as above.

### Chromatin associated poly(A)-RNA enrichment and LC-MS/MS

Poly(A)-RNA isolation was performed with modifications to a previously described protocol (Iadevaia et al. 2018). A549 cells were grown in the presence of 0.01% DMSO, 1 µM GSK591 (Cayman), or 1 µM MS023 (Cayman) for two days. Cells were washed with 4 °C PBS and irradiated on ice with 100 mJ cm^-2^ in a UV Stratalinker 1800. Cells were centrifuged at 500 x g for 10 minutes at 4 °C. Chromatin was isolated with nuclear lysis buffer (NLB; 10 mM Tris-HCl pH 7.5 at 4°C, 0.1% NP-40, 400 mM KCl, 1 mM DTT) supplemented with 40 U/mL RNaseOUT (Thermo), protease inhibitor (Thermo), and phosphatase inhibitor (Thermo). The chromatin pellet was resuspended in NLB and sonicated for five seconds at 20% amplitude with a probe-tip sonicator using a 1/8” tip. The sonicate was centrifuged at 10,000 x g for 10 minutes and the soluble material transferred to a low-adhesion RNase-free microcentrifuge tube. An aliquot from each sample was saved to serve as the unenriched control. The samples were split into two separate tubes, one of which received 10 µg of competitor 25-nt poly(A) RNA. Magnetic oligo- (dT) beads (NEB) were equilibrated in NLB and added to the enrichments. The samples were vortexed at room temperature for 10 minutes. The beads were then captured on a magnetic column, and the supernatant transferred to fresh tube for additional rounds of depletion. The beads were washed once with buffer A (10 mM Tris pH 7.5, 600 mM KCl, 1 mM EDTA, 0.1% Triton X-100), followed by buffer B (10 mM Tris pH 7.5, 600 mM KCl, 1 mM EDTA) and lastly buffer C (10 mM Tris pH 7.5, 200 mM KCl, 1 mM EDTA). The RNA was eluted by incubating the beads in 10 µL of 10 mM Tris pH 7.5 at 80 °C for two minutes, capture of magnetic beads using a magnetic column, and quickly transferring the supernatant to a new tube. The beads were then used for two additional rounds of poly(A)-RNA capture.

The (un)enriched proteome were treated with Benzonase (Sigma) and then digested with trypsin (Promega) prior to performing LC-MS/MS. Proteins were resuspended in 5% SDS and 5 mM DTT and left incubating for 1 hour at room temperature. Samples were then alkylated with 20 mM iodoacetamide in the dark for 30 minutes. Afterward, phosphoric acid was added to the sample at a final concentration of 1.2%. Samples were diluted in six volumes of binding buffer (90% methanol and 10 mM ammonium bicarbonate, pH 8.0). After gentle mixing, the protein solution was loaded to an S-trap filter (Protifi) 96-well plate and spun at 500 x g for 30 sec. The sample was washed twice with binding buffer. Finally, 100 ng of sequencing grade trypsin (Promega), diluted in 50 mM ammonium bicarbonate, was added into the S-trap filter and samples were digested at 37 °C for 18 hours. Peptides were eluted in three steps: (i) 40 µl of 50 mM ammonium bicarbonate, (ii) 40 µl of 0.1% TFA and (iii) 40 µl of 60% acetonitrile and 0.1% TFA. The peptide solution was pooled, spun at 1,000 x g for 30 sec and dried in a vacuum centrifuge. Prior to mass spectrometry analysis, samples were desalted using a 96-well plate filter (Orochem) packed with 1 mg of Oasis HLB C-18 resin (Waters). Briefly, the samples were resuspended in 100 µL of 0.1% TFA and loaded onto the HLB resin, which was previously equilibrated using 100 µL of the same buffer. After washing with 100 µL of 0.1% TFA, the samples were eluted with a buffer containing 70 µL of 60% acetonitrile and 0.1% TFA and then dried in a vacuum centrifuge. For LC-MS/MS acquisition, samples were resuspended in 10 µL of 0.1% TFA and loaded onto a Dionex RSLC Ultimate 300 (Thermo Scientific), coupled online with an Orbitrap Fusion Lumos (Thermo Scientific). Chromatographic separation was performed with a two-column system, consisting of a C-18 trap cartridge (300 µm ID, 5 mm length) and a picofrit analytical column (75 µm ID, 25 cm length) packed in-house with reversed-phase Repro-Sil Pur C18-AQ 3 µm resin. Peptides were separated using a 60-minute gradient from 4-30% buffer B (buffer A: 0.1% formic acid, buffer B: 80% acetonitrile + 0.1% formic acid) at a flow rate of 300 nL/min. The mass spectrometer was set to acquire spectra in a data-dependent acquisition (DDA) mode. The full MS scan was set to 300-1200 m/z in the orbitrap with a resolution of 120,000 (at 200 m/z) and an AGC target of 5 x 10^5^. MS/MS was performed in the ion trap using the top speed mode (2 secs), an AGC target of 10^4^ and an HCD collision energy of 35. Raw files were searched using Proteome Discoverer software (v2.4, Thermo Scientific) using SEQUEST search engine with the SwissProt human database. The search for total proteome included variable modification of N-terminal acetylation and fixed modification of carbamidomethyl cysteine. Trypsin was specified as the digestive enzyme with up to 2 missed cleavages allowed. Mass tolerance was set to 10 pm for precursor ions and 0.2 Da for product ions. Peptide and protein false discovery rate was set to 1%.

Data were transformed, normalized and statistics were applied as described previously (Aguilan et al. 2020). Briefly, the data were log_2_ transformed and normalized by the average of the data distribution. Statistical analysis was performed by using a two-tail heteroscedastic t-test.

### Immunoprecipitation

A549 cells were grown with 0.01% DMSO, 1 µM GSK591 (Cayman), or 1 µM MS023 (Cayman) for two days. Cells were harvested using trypsin (Corning) and washed once with 4 °C PBS supplemented with PRMTi. Cells were then resuspended in RIPA buffer (1% NP-40, 150 mM NaCl, 50 mM Tris-HCl pH 8 at 4 °C, 0.25% sodium deoxycholate, 0.1% SDS, and 1 mM EDTA) supplemented with 40 U/mL RNaseOUT (Thermo) and protease inhibitor (Thermo). Lysates were incubated on ice for 10 minutes followed by sonication for five seconds at 20% amplitude with a probe-tip sonicator using a 1/8” tip. Lysates were then spun at 10,000 x g for 10 minutes at 4°C. Supernatants were transferred to new low-adhesion RNase-free microcentrifuge tubes and normalized to the same protein concentration using bicinchoninic acid (Pierce). Primary antibody targeting CHTOP (LSBio, LS-C193506) or SNRPB (ProteinTech, 16807-1-AP) was added followed by incubation overnight at 4°C with gentle rotation. The next morning, Protein G agarose (Millipore-Sigma 16-201) was equilibrated in lysis buffer and added to the lysates at 4 °C with gentle rotation. The beads were washed three times with lysis buffer followed by resuspension in 1x Laemmli buffer for western blotting with CHTOP (as above), SNRPB (as above), Rme2s (CST, 13222), and Rme2a (CST, 13522) antibodies.

### CRISPRi and expression of methylarginine mutants

The dCas9-KRAB-MeCP2 expression vector (VB900120-5303pyt) was purchased from VectorBuilder (Santa Clara, CA). The PRMT-targeting gRNA parent vector (VB210119-1169qwd) was purchased from VectorBuilder and custom gRNA sequences (**Supplemental Table 2**) derived from CRISPick (Doench et al. 2016) were cloned as described previously (Ran et al. 2013) with the exception that BfuA1 (NEB; R0701S) was used to digest the parent vector. Sm wildtype (VB210103-1026dg), R-to-A (VB210103-1027ttz), and R-to-K (VB210317-1185yxr) as well as CHTOP wildtype (VB210427-1238rhx), R-to-A (VB210427-1241jrw), and R-to-K (VB210427-1242gjh) expression vectors were cloned by VectorBuilder. Lentiviral particles containing expression vectors were produced in HEK293T cells using calcium phosphate transfection. Transduction of A549 cells was accomplished by combining lentivirus with polybrene containing media (4 µg/mL) and centrifuging at 30 °C for 90 minutes at 500 x g followed by incubation for 24 hours at 37 °C, 5% CO_2_ with humidity. Following the 24-hour incubation, lentiviral containing media was removed and complete DMEM containing appropriate selection antibiotic (Cayman) was added. A549 cells expressing dCas9-KRAB-MeCP2 were selected using 2 µg/mL puromycin and then maintained in culture with 1 µg/mL puromycin. For PRMT knockdown and Sm expression experiments, A549 cells were selected with 10 µg/mL blasticidin for 72 hours. For CHTOP expression experiments, A549 cells were selected for 96 hours with 1 mg/mL G418. Primers used in RT-qPCR experiments can be found in **Supplemental Table 2**. Lysis for western blotting was accomplished as above with FLAG (Sigma, F1804), Rme2s (as above), and GAPDH (Abcam, ab9484) antibodies.

### Cell fractionation

A549 cells were grown with 0.01% DMSO, 1 µM GSK591 (Cayman), or 1 µM MS023 (Cayman) for two days. Cells were harvested using trypsin (Corning) and washed once with 4 °C PBS supplemented with PRMTi. Cells were resuspended in Hypotonic Lysis Buffer (10 mM TrisCl pH 8, 0.1% NP-40, 1 mM KCl, 1.5 mM MgCl_2_, supplemented with protease inhibitor and 40 U/mL RNaseOUT) and rotated for 30 minutes at 4 °C. Cells were then centrifuged at 10,000 x g for 10 minutes at 4 °C and the supernatant was kept as the cytosolic fraction. The nuclear pellet was washed once with hypotonic buffer to reduce carry over then resuspendned in NLB (as above) and rotated for 30 minutes at 4 °C. After centrifugation at 10,000 x g for 10 minutes the supernatant was kept as the nucleoplasm and the remaining material was washed once with NLB and kept as the chromatin fraction. RNA from each fraction was isolated using TRIzol (Thermo) and RT-qPCR was performed as above. For western analysis, the chromatin fraction was resuspended in NLB then sheared with sonication. This was followed by addition of Laemmli buffer to 1x and blotting with H3 (Abcam, ab1791), CHTOP, and SNRPB antibodies (as above).

### Statistical analysis

All western blots were performed independently at least twice. RT-qPCR was performed at least three-times with independent biological replicates. Statistical analyses were performed using either Prism software (Version 8.3.1, GraphPad) or R (version 4.0.2). To compare general distributions, the Kolmogorov-Smirnov test was used. To compare median distribution changes the Wilcoxon rank-sum test was used. To account for differences in sample size between global and RI distributions, random sampling from the global population equivalent to the number of RI within the tested condition was performed. This process was repeated one thousand times after which the median *P* was reported. To compare means where only two groups exist, independent t-test was performed. To compare means where two categorical variables exist, two-way ANOVA with post-hoc Dunnett’s test was performed. GeneOverlap with Fisher’s exact test was used to determine odds ratios of RI overlap between different RNA-seq datasets (Shen 2020). All code used to generate data in this manuscript can be found here: https://github.com/Shechterlab/PRMTsRegulatePostTranscriptionalDI.

## ACKNOWLEDGMENTS

This work was supported by the National Institutes of Health [R01GM108646 to D.S., GM57829 to C.C.Q., R01GM134379 to M.J.G], the American Lung Association Discovery Award LCD-564723 [D.S] and the HIRSCHL/MONIQUE WEILL-CAULIER TRUSTS to M.J.G and also to D.S.. We thank Dr. Martin Krzywinski (http://mkweb.bcgsc.ca/) for help with data visualization.

## AUTHOR CONTRIBUTIONS

M.I.M. conceived and designed the project, performed biochemical and bioinformatic experiments, interpreted resulting data, and wrote the manuscript. A.D.C. performed SKaTER-seq analysis. V.G. performed bioinformatic analysis. S.S. performed LC-MS/MS analysis. M.J.G. and C.C.Q. helped with experimental design and data interpretation. D.S. supervised the study and helped with experimental design, data interpretation, and manuscript writing. All authors reviewed the manuscript.

## ACCESSION NUMBERS

Raw data for RNA seq and SKaTER seq is deposited under GEO (GSExxxxxx). Accessions for publicly available RNA-seq data used in this study can be found in **Supplemental Table 3**. Raw data for chromatin-associated poly(A) LC-MS/MS is deposited under Chorus (xxxx).

## SUPPLEMENTAL FIGURE LEGENDS

**Supplemental Figure 1.**
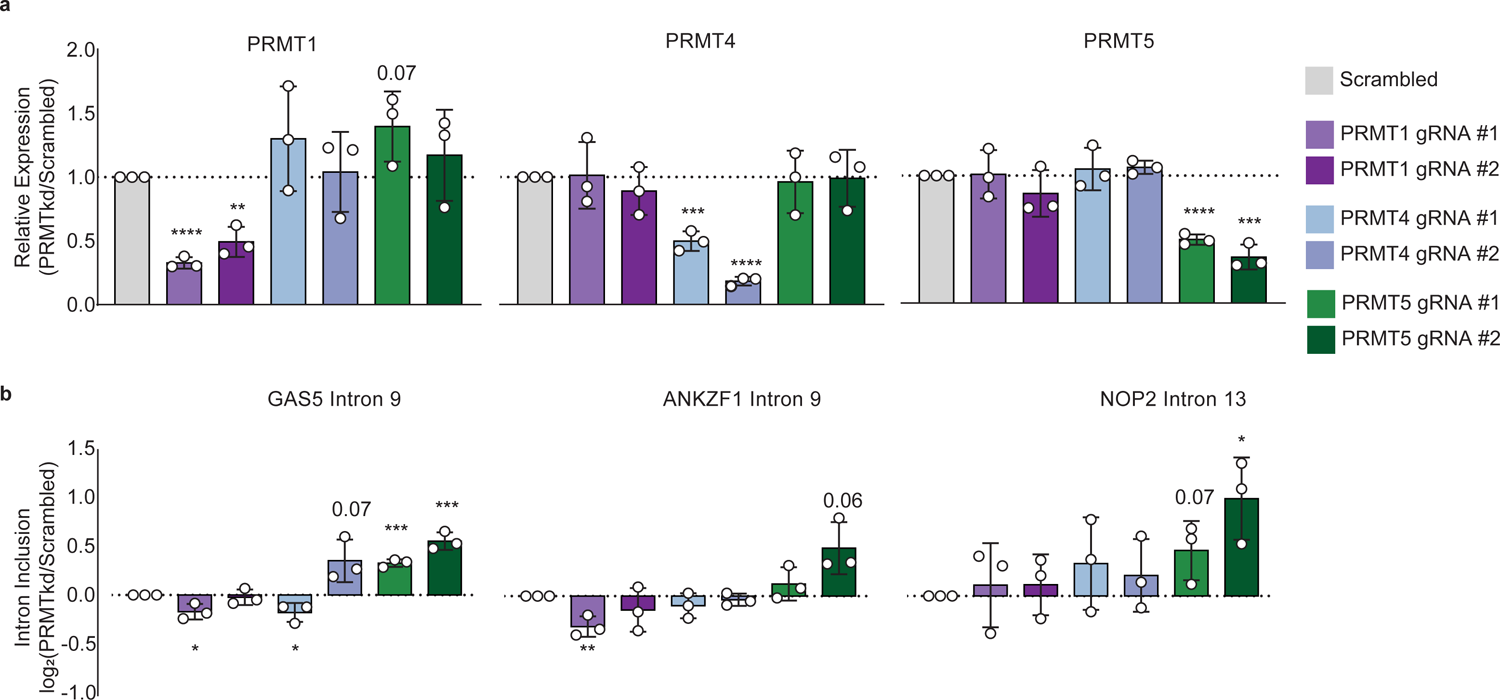
PRMTkd recapitulates RI inclusion seen with PRMTi. **a-b.** RT-qPCR of PRMT expression **(a)** or RI **(b)** 96 hours after transduction of A549 cells with gRNAs targeting the indicated PRMT. Data are represented as mean ± SD. Significance determined using Student’s t-test; * < 0.05, ** < 0.01, *** < 0.001, **** < 0.0001.

**Supplemental Figure 2.**
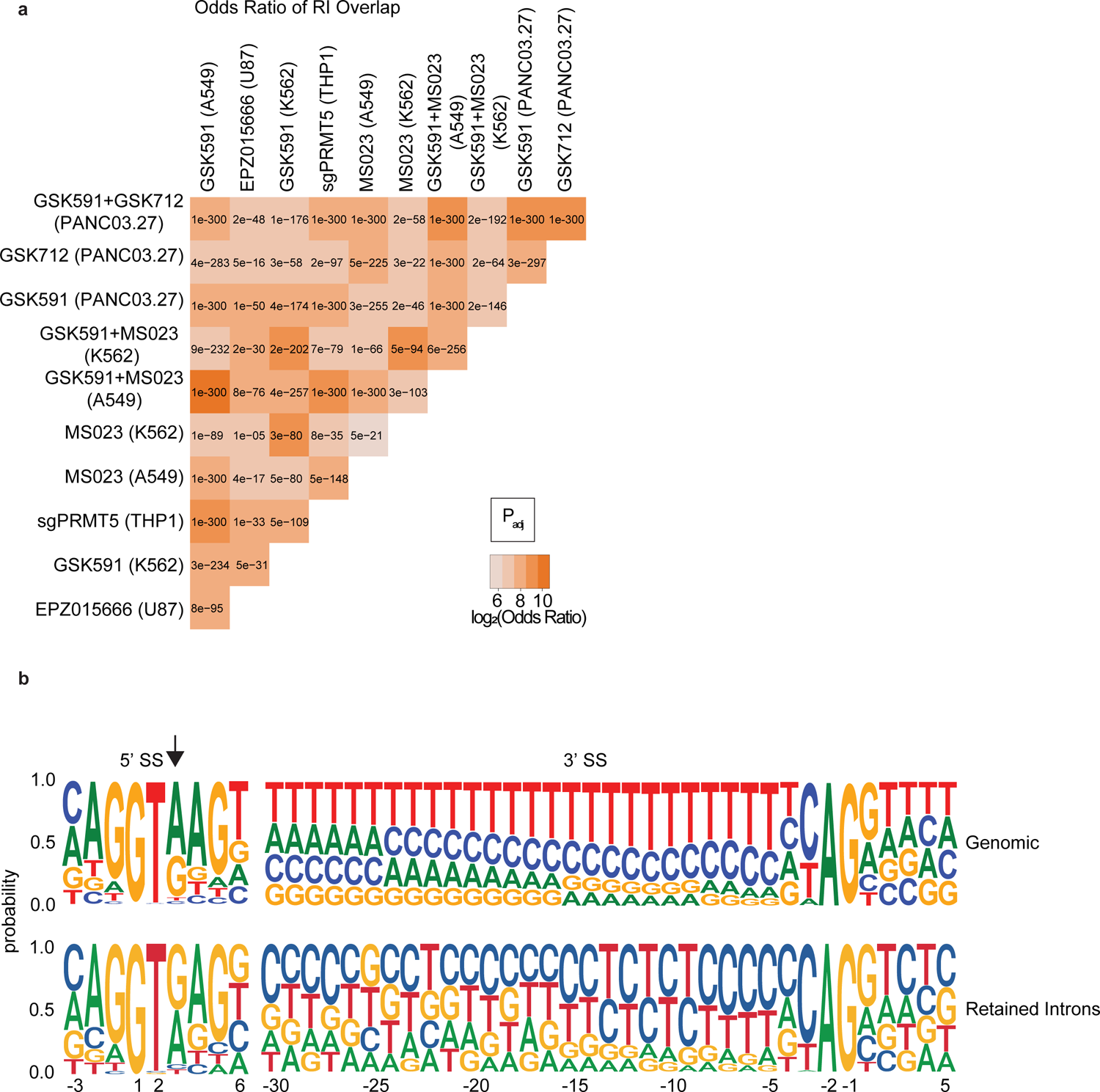
PRMTs regulate a conserved class of RI that share unique features. **a.** Matrix comparing the log_2_(odds ratio) and significance as determined by the Fisher’s Exact Test of overlapping RI from indicated experimental models. **b.** Web logo diagram of nucleotide distribution probability of A549 expressed or retained introns.

**Supplemental Figure 3.**
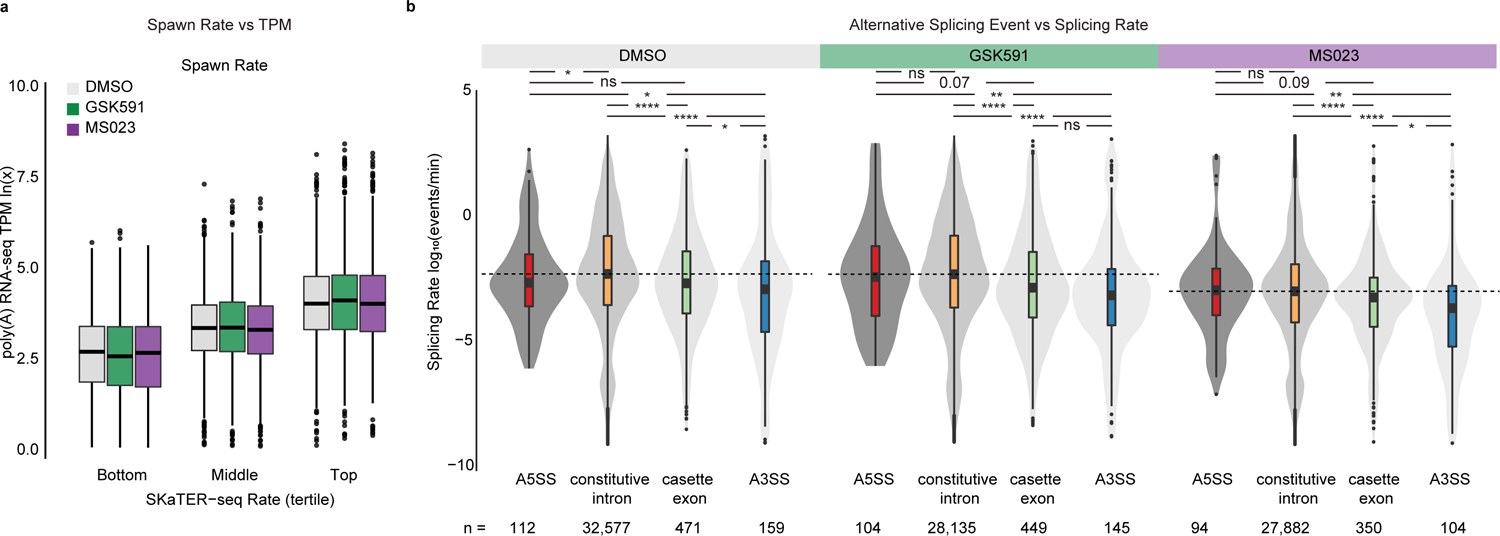
Transcript spawn rate correlates with expression and alternative splicing events are slower than constitutive ones. **a.** Correlation of RNA pol II initiation and pause-release (spawn rate) with poly(A)-RNA seq transcripts per million (TPM). x-axis is spawn rate in tertiles; y-axis is poly(A)-RNA seq ln(TPM). **b.** Distribution of splicing rate versus splicing event as determined by SKaTER-seq modeling. A5SS = alternative 5’ splice site, A3SS = alternative 3’ splice site. Dashed line indicates constitutive intron median; solid line within boxplot is event-specific median. Significance determined using Kolmogorov-Smirnov test; * < 0.05, ** < 0.01, *** < 0.001, **** < 0.0001, ns = not significant.

**Supplemental Figure 4.**
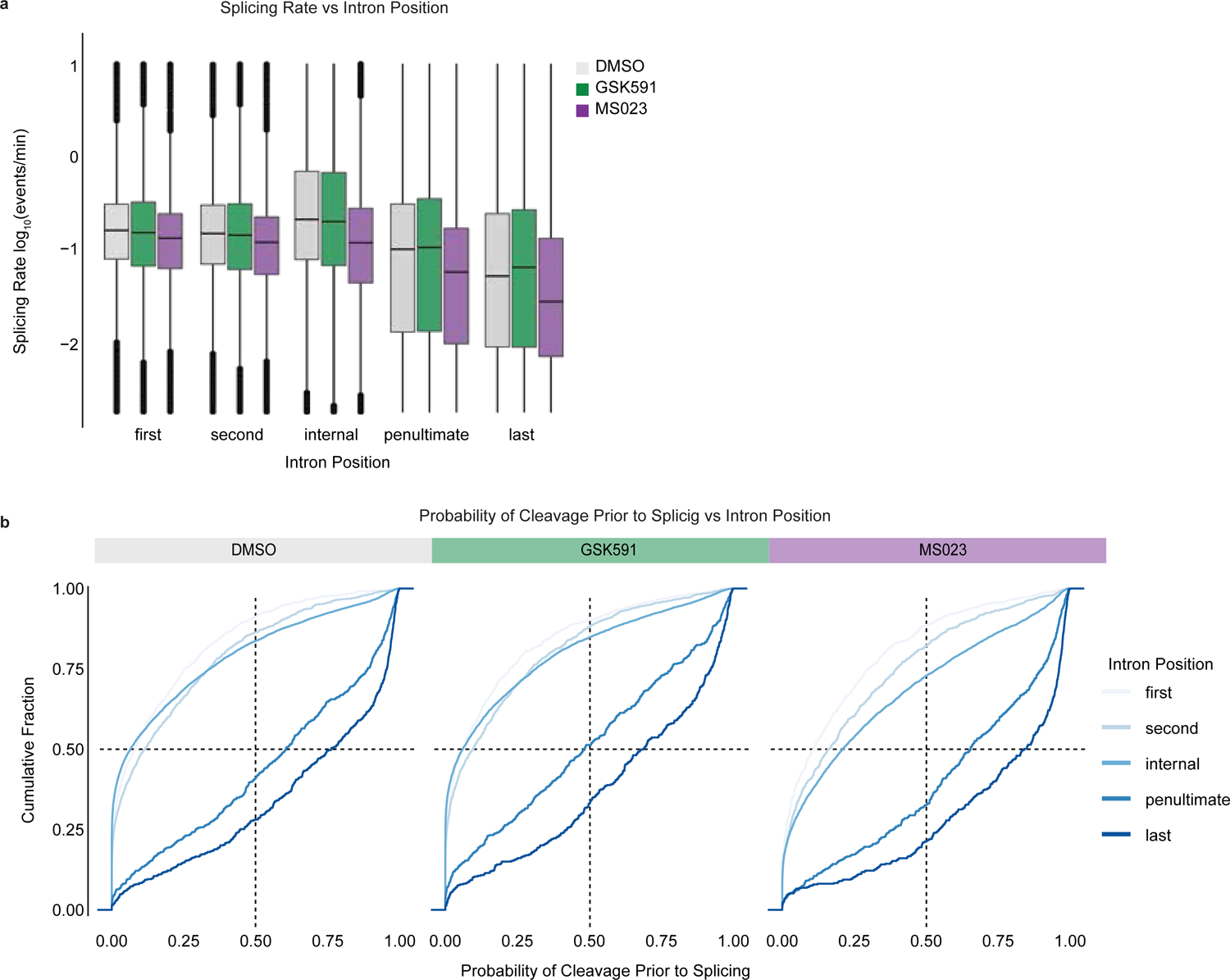
Splicing rate and probability of cleavage prior to splicing correlate with intron position. **a.** Correlation of splicing rate and intron position in cells treated with DMSO, GSK591, or MS023 for two-days. **b.** Cumulative distribution functions comparing the probability of transcript cleavage prior to intron splicing and intron position. Color range indicates intron position where light blue is closer to transcription start site and dark blue closer to transcription end site.

**Supplemental Figure 5.**
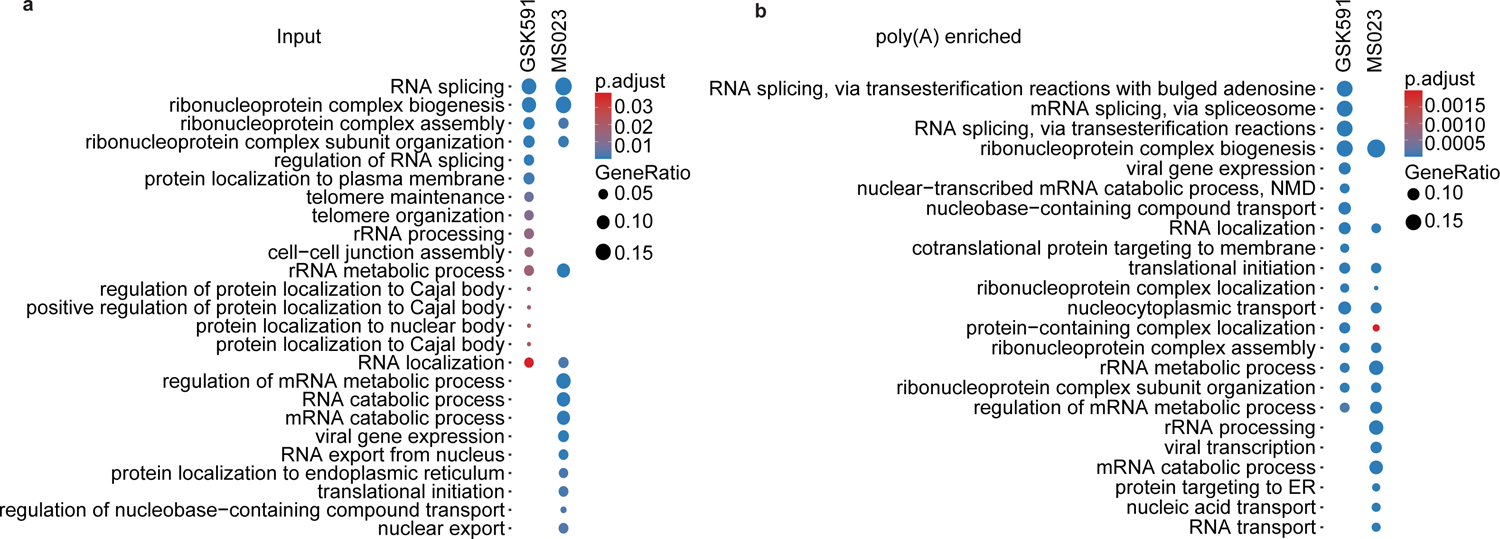
PRMTi alters association of RNA splicing, localization, and processing factors to chromatin and chromatin-associated poly(A) RNA. **a-b.** Enriched biological processes found in input chromatin **(a)** and chromatin-associated poly(A) RNA **(b)** in A549 cells treated with GSK591 or MS023 relative to DMSO.

**Supplemental Figure 6.**
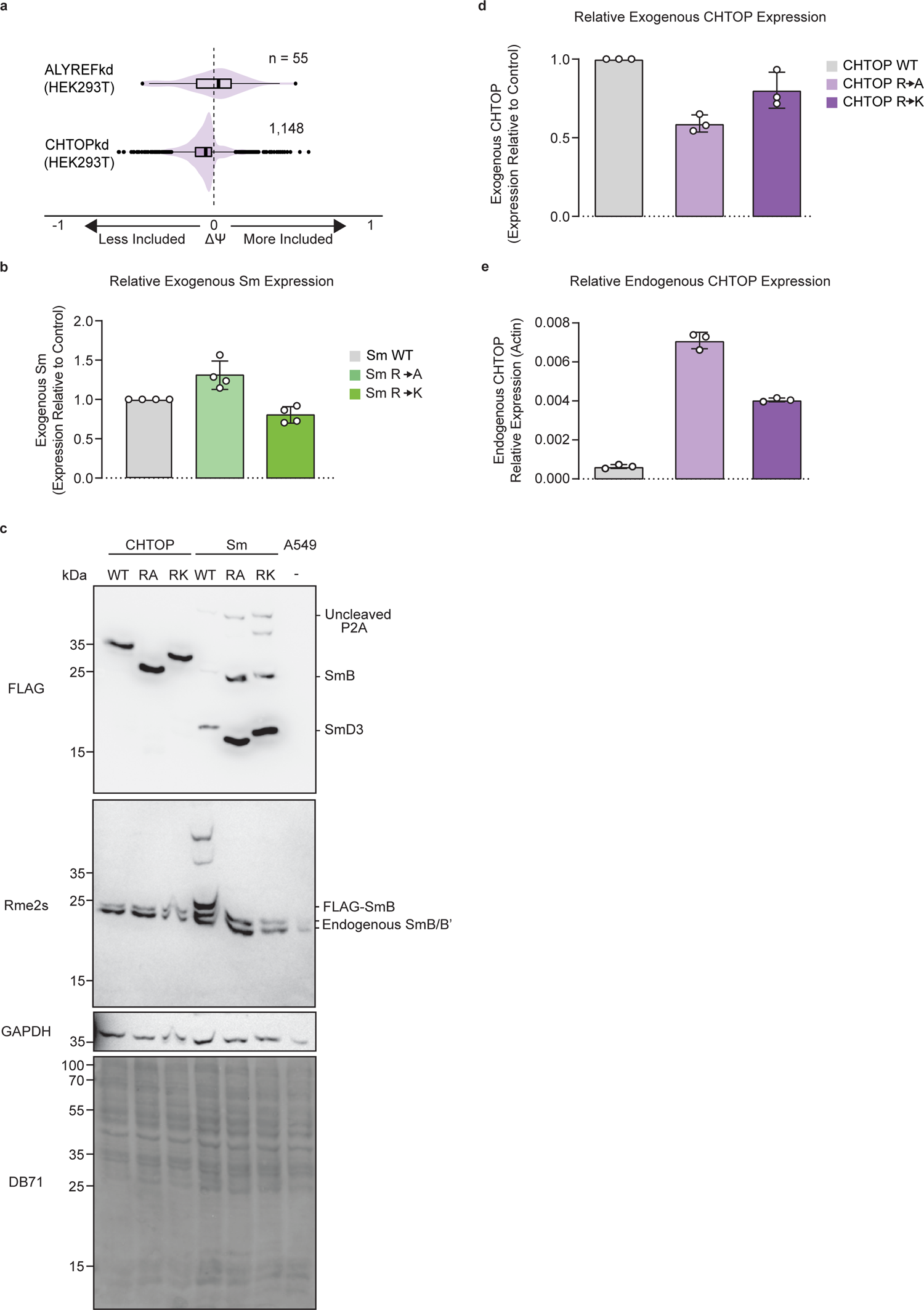
CHTOP and Sm arginine mutants phenocopy PRMTi. **a.** Comparison of ΔΨ for RI following ALYREFkd and CHTOPkd where ΔΨ = Ψ (Treatment) – Ψ (Control). **b.** RT-qPCR of Exogenous Sm R-to-A and R-to-K mutants normalized to wildtype expression vector. Data are represented as mean ± SD. **c.** Western blot of total cell extract from A549 cells expressing CHTOP or Sm wildtype, R-to-A, or R-to-K mutants. **d.** RT-qPCR of Exogenous CHTOP R-to-A and R-to-K mutants normalized to wildtype expression vector. Data are represented as mean ± SD. **e.** RT-qPCR of Endogenous CHTOP in A549 cells expressing CHTOP wildtype, R-to-A, or R-to-K mutants. Data are represented as mean ± SD.

---

Supplemental Table 1 – Chromatin-Associated Poly(A) RNA Enrichment Enrichment and relative abundance of proteins in chromatin or chromatin-associated poly(A) RNA following treatment with DMSO, GSK591, or MS023 for two days.

Supplemental Table 2 – Primer Sequences for RT-qPCR and gRNAs

Supplemental Table 3 – Publicly available data used in this study

